# Optimal policy for attention-modulated decisions explains human fixation behavior

**DOI:** 10.1101/2020.08.04.237057

**Authors:** Anthony Jang, Ravi Sharma, Jan Drugowitsch

## Abstract

Traditional accumulation-to-bound decision-making models assume that all choice options are processed simultaneously with equal attention. In real life decisions, however, humans tend to alternate their visual fixation between individual items in order to efficiently gather relevant information [46, 23, 21, 12, 15]. These fixations also causally affect one’s choices, biasing them toward the longer-fixated item [38, 2, 25]. We derive a normative decision-making model in which fixating a choice item boosts information about that item. In contrast to previous models [25, 39], we assume that attention enhances the reliability of information rather than its magnitude, consistent with neurophysiological findings [3, 13, 29, 45]. Furthermore, our model actively controls fixation changes to optimize information gathering. We show that the optimal model reproduces fixation patterns and fixation-related choice biases seen in human decision-makers, and provides a Bayesian computational rationale for the fixation bias. This insight led to additional behavioral predictions that we confirmed in human behavioral data. Finally, we explore the consequences of changing the relative allocation of cognitive resources to the attended versus the unattended item, and show that decision performance is benefited by a more balanced spread of cognitive resources.

## Introduction

Would you rather have a donut or an apple as a mid-afternoon snack? If we instantaneously knew their associated rewards, we could immediately choose the higher-rewarding option. However, such decisions usually take time and are variable, suggesting that they arise from a neural computation that extends over time [33, 37]. If we assume these computations to involve a stream of noisy samples of each item’s underlying value, then, normatively, we would accumulate these samples over time until we can confidently choose the higher-rewarding item [42]. This strategy can be implemented with accumulation-to-bound models which show that choices between equally desirable items are slower and more variable than choices between items with a larger difference in desirability, as is the case in human behavior [34].

Standard accumulation-to-bound models assume that all choice options receive equal attention during decision-making. However, the ability to drive one’s attention amidst multiple, simultaneous trains of internal and external stimuli is an integral aspect of everyday life. Indeed, humans tend to alternate between fixating on different items when making decisions. Furthermore, their final choices are biased towards the item that they looked at longer, irrespective of its desirability [38, 25, 26, 11]. This choice bias has been previously replicated with a modified accumulation-to-bound model. However, this model assumed that fixations are driven by brain processes that are exogenous to the computations involved in decision-making [25]. This stands in contrast to studies of visual attention, where fixations appear to be controlled to extract choice-relevant information in a statistically efficient manner. In this case, fixations are driven by processes endogenous to the decision [46, 23, 21, 12, 15].

We asked if the choice bias associated with fixations can be explained with a unified framework in which fixation changes and decision-making are part of the same process. To do so, we endowed normative decision-making models [42] with attention that boost the amount of information one collects about each choice option, in line with neurophysiological findings [3, 13, 29, 45]. We furthermore assumed that this attention was overt [32, 20], and thus reflected in the decision maker’s gaze which was controlled by the decision-making process.

We show that, under these assumptions, the normative decision-making strategy featured the same choice bias as observed in human decision makers: it switched attention more frequently when deciding between items with similar values, and was biased towards choosing items that were attended last, and attended longer. It furthermore led to new predictions that we could confirm in human behavior: choice biases varied based on the amount of time spent on the decision and the average desirability across both choice items. Lastly, it revealed why the observed choice biases might, in fact, be rational. Overall, our work provides a unified framework in which the optimal, attention-modulated information-seeking strategy naturally leads to biases in choice that are driven by visual fixations, as observed in human decisions.

## Results

### An attention-modulated decision-making model

Before describing our attention-modulated decision-making model, we will first briefly recap the attention-free model [42] that ours builds upon. This model assumes that for each decision trial, a true value associated with each item (*z*_1_, *z*_2_) is drawn from a normal prior distribution with mean 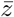 and variance 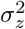. Therefore, 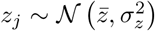 for both *j* ∈ {1, 2}. We assume the decision maker knows the shape of the prior, but can’t directly observe the drawn true values. In other words, the decision maker a-priori knows the range of values associated with the items they need to compare, but doesn’t know what exact items to expect nor what their associated rewards will be. For example, one such draw might result in a donut and an apple, each of which has an associated value to the decision maker (i.e., satisfaction upon eating it). In each *n*th time step of length *δt*, they observe noisy samples centered around the true values, called *momentary evidence*, 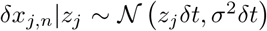. While the model is agnostic to the origin of these samples, they might arise from computations to infer the items’ values (e.g., how much do I currently value the apple?), memory recall (e.g., how much did I previously value the apple?), or a combination thereof [37]. As the decision maker’s aim is to choose the higher-valued item, they ought to accumulate evidence for some time to refine their belief in the items’ values. Once they have accumulated evidence for *t* = *Nδt* seconds, their posterior belief for the value associated with either item is

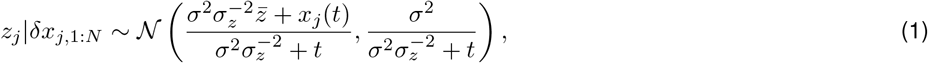

where 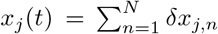 is the accumulated evidence for item j [42]. The variance of this posterior reflects the uncertainty in the decision maker’s value inference. In the attention-free model, this uncertainty monotonically decreases identically over time for both items, reflecting the standard assumption of accumulation-to-bound models that, in each small time period, the same amount of evidence is gathered for either choice item.

To introduce attention-modulation, we assume that attention limits information about the unattended item (Fig. 1A,B). This is consistent with behavioral and neurophysiological findings showing that attention boosts behavioral performance [13, 14, 44] and the information encoded in neural populations [31, 36, 45]. To limit information, we change the momentary evidence for the unattended item *j* to 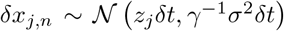, while leaving that for the attended item unchanged. Here, the *γ* term controls the degree to which inattention leads to noisier acquisition of information (0 < *γ* ≤ 1). Setting γ = 1 recovers the attention-free scenario [42].

**Figure 1:**
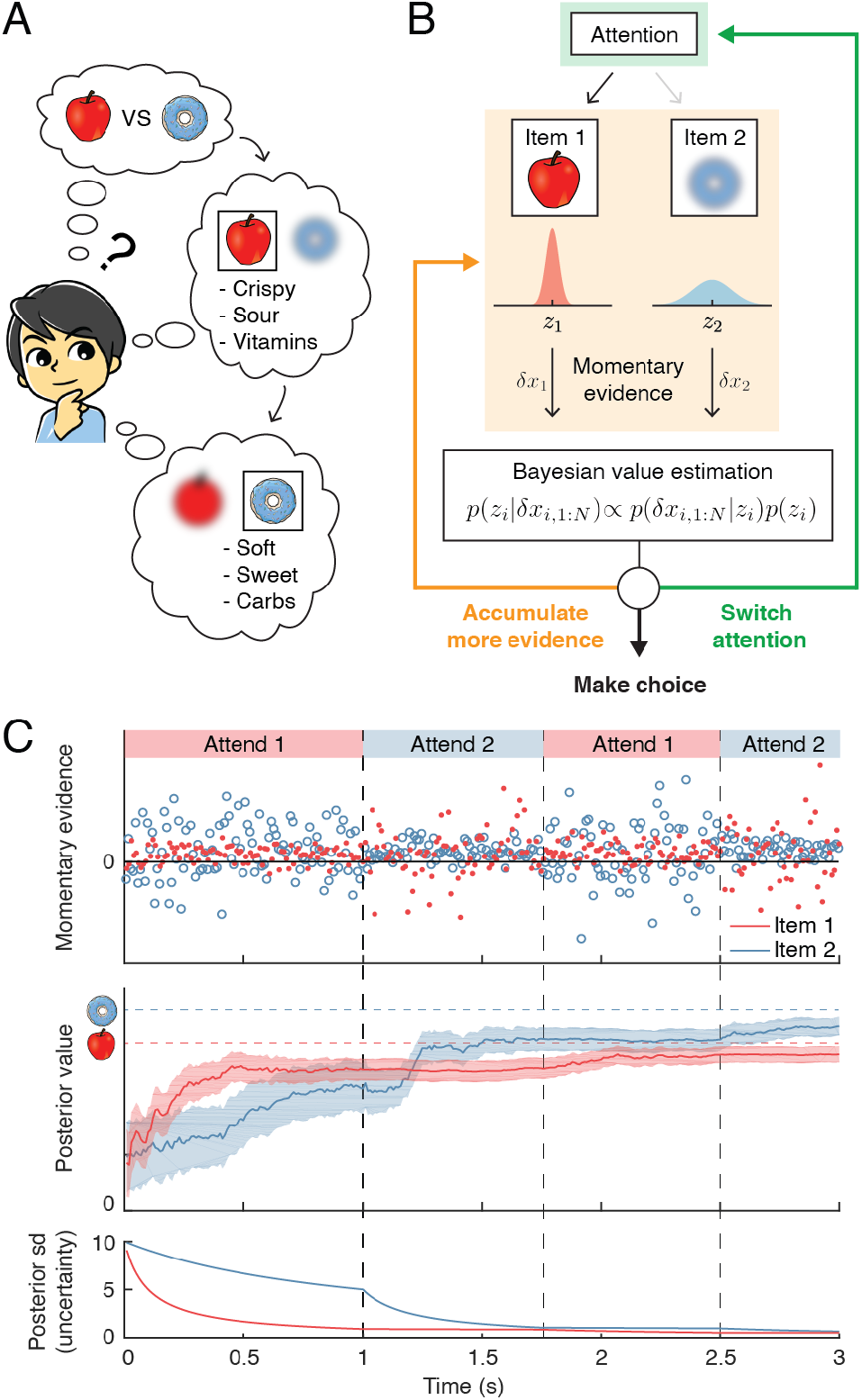
Attention-modulated evidence accumulation. (A) Schematic depicting the value-based decision-making model. When choosing between two snack items (e.g., apple versus donut), people tend to evaluate each item in turn, rather than think about all items simultaneously. While evaluating one item, they will pay less attention to the unattended item (blurred item). (B) Schematic of the value-based decision process for a single trial. At trial onset, the model randomly attends to one item (green box). At every time step, it accumulates momentary evidence (orange box) that provides information about the true value of each item, which is combined with the prior belief of each item’s value to generate a posterior belief. Note that the momentary evidence of the attended item comes from a tighter distribution. Afterwards, the model assesses whether to accumulate more evidence (orange), make a choice (black), or switch attention to the other item (green). (C) Evolution of the evidence accumulation process. The top panel shows momentary evidence at every time point for the two items. Note that evidence for the unattended item has a wider variance. The middle panel shows how the posterior estimate of each item may evolve over time (mean ± 1SD). The dotted lines indicate the unobserved, true values of the two items. The bottom panel shows how uncertainty decreases regarding the true value of each item. As expected, uncertainty decreases faster for the currently attended item compared to the unattended one.

Lowering information for the unattended item impacts the value posteriors as follows. If the decision maker again accumulates evidence for some time *t* = *Nδt*, their belief about item *j* = 1’s value changes from Eq. (1) to

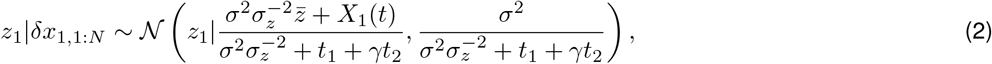

where *t*_1_ and *t*_2_, which sum up to the total accumulation time (*t* = *t*_1_ + *t*_2_), are the durations that items 1 and 2 have been attended, respectively. The accumulated evidence *X*_1_(*t*) now isn’t simply the sum of all momentary pieces of evidence, but instead down-weights them by *γ* if the associated item is unattended (see Methods). This prevents the large inattention noise from swamping the overall estimate [16]. An analogous expression provides the posterior *z*_2_|*δx*_2,1:*N*_ for item 2 (Supplementary Information).

The attention modulation of information is clearly observable in the variance of the value’s posterior (Eq. (2)). For *γ* < 1, this variance, which is proportional to the decision maker’s uncertainty about the option’s value, drops more quickly over time if item 1 rather than item 2 is attended (i.e., if *t*_1_ rather than *t*_2_ increases). Therefore, it depends on how long each of the two items have been attended to, and might differ between the two items across time (Fig. 1C). As a result, decision performance depends on how much time is allocated to attending to each item.

The decision maker’s best choice at any point in time is to choose the item with the larger expected value, as determined by the value posterior. However, the posterior by itself does not determine when it is best to stop accumulating evidence. In our previous attention-free model, we addressed the optimal stopping time by assuming that accumulating evidence comes at cost c per second, and found the optimal decision policy under this assumption [42]. Specifically, at each time step of the decision-making process, the decision maker could choose between three possible actions. The first two actions involve immediately choosing one of the two items, which promises the associated expected rewards. The third action is to accumulate more evidence that promises more evidence, better choices, and higher expected reward, but comes at a higher cost for accumulating evidence. We found the optimal policy using dynamic programming that solves this arbitration by constructing a value function that, for each stage of the decision process, returns all expected rewards and costs from that stage onward [6, 8]. The associated policy could then be mechanistically implemented by an accumulation-to-bound model that accumulates the difference in expected rewards, Δ = 〈*z*_2_|*δx*_2,1:*N*_〉 – 〈*z*_1_|*δx*_1,1:*N*_), and triggers a choice once one of two decision boundaries, which collapse over time, is reached [42].

Once we introduce attention, a fourth action becomes available: the decision maker can choose to switch attention to the currently unattended item (Fig. 1B). If such a switch comes at no cost, then the optimal strategy would be to continuously switch attention between both items to sample them evenly across time. We avoid this physically unrealistic scenario by introducing a cost *c_s_* for switching attention. This cost also includes a switch time *t_s_*, which is not included in *t*_1_ and *t*_2_. Overall, this leads to a value function defined over a four-dimensional space: the expected reward difference Δ, the evidence accumulation times *t*_1_ and *t*_2_, and the currently attended item *y* ∈ {1, 2} (see Supplementary Information). As the last dimension can only take one of two values, we can equally use two three-dimensional value functions. This results in two associated policies that span the three-dimensional *state space* (Δ, *t*_1_, *t*_2_) (Fig. 2).

**Figure 2:**
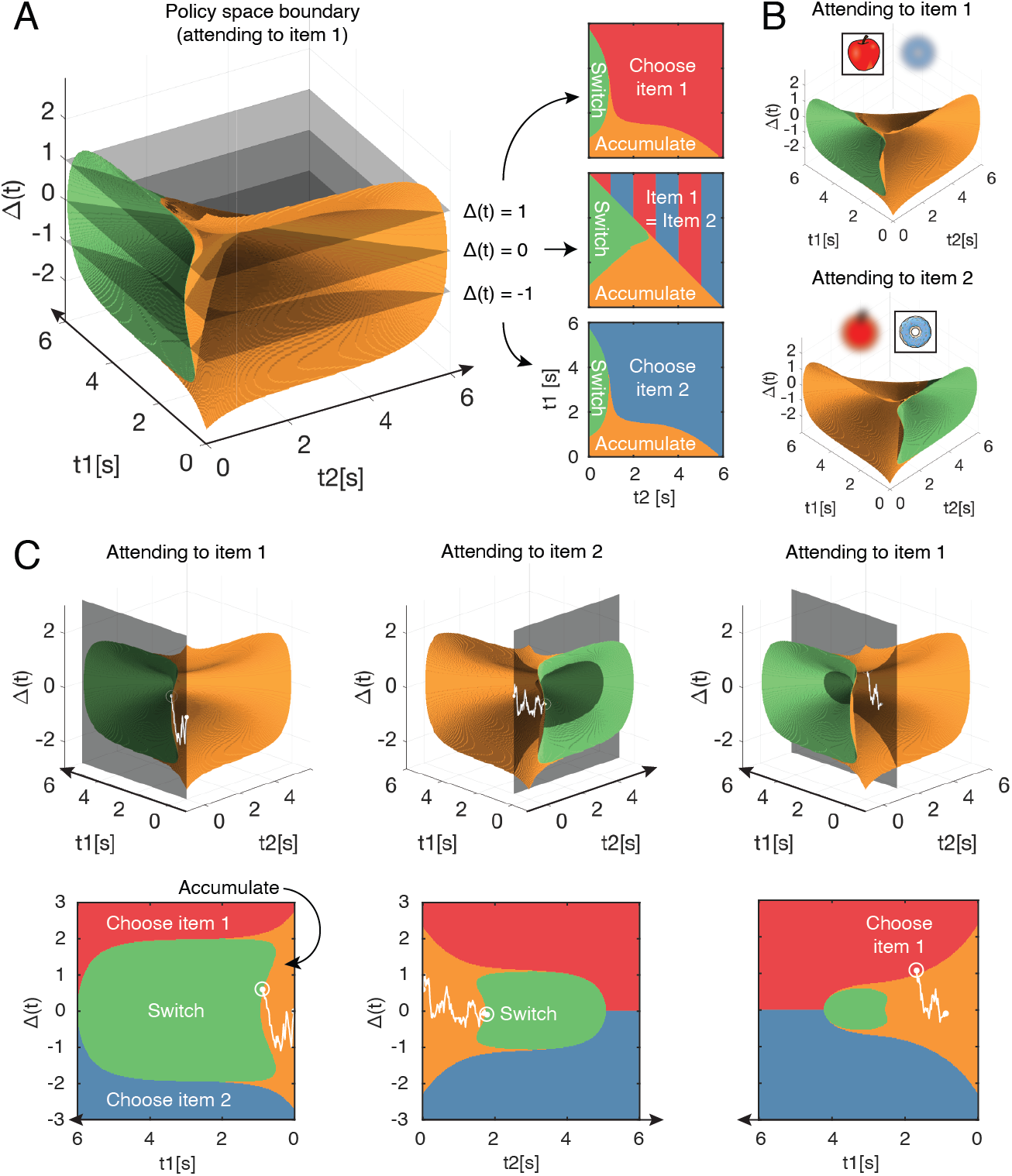
Navigating the optimal policy space. (A) The optimal policy space. The policy space can be divided into regions associated with different optimal actions (choose item 1 or 2, accumulate more evidence, switch attention). The boundaries between these regions can be visualized as contours in this space. The three panels on the right show cross-sections after slicing the space at different Δ values. Note that when Δ = 0 (middle panel), the two items have equal value and therefore there is no preference for one item over the other. (B) Optimal policy spaces for different values of *y* (currently attended item). The two policy spaces are mirror-images of each other. (C) Example deliberation process of a single trial demonstrated by a particle that diffuses across the optimal policy space. In this example, the model starts by attending to item 1, then makes two switches in attention before eventually choosing item 1. The bottom row shows the plane in which the particle diffuses. Note that the particle diffuses on the (grey, shaded) plane perpendicular to the time axis of the unattended item, such that it only increases in *t*_j_ when attending to item j. Also note that the policy space changes according to the item being attended to, as seen in (B). See results text for more detailed description.

### Features of the optimal policy

At any point within a decision, the modelâĂŹs current state is represented by a location in this 3D policy space, such that different regions in this space designate the optimal action to perform (i.e., choose, accumulate, switch). The boundaries between these regions can be visualized as contours in this 3D state space (Fig. 2A). As previously discussed, there are two distinct policy spaces for when the decision maker is attending to item 1 versus item 2 that are symmetric to each other (Fig. 2B).

Within a given decision, the deliberation process can be thought of as a particle that drifts and diffuses in this state space. The model starts out attending to an item at random (*y* ∈ 1,2), which determines the initial policy space (Fig. 2B). Assume an example trial where the model attends to item 1 initially (*y* = 1). At trial onset, the decision maker holds the prior belief, such that the particle starts on the origin (Δ = 0, *t*_1_ = *t*_2_ = 0) which is within the âĂIJaccumulateâĂi region. As the model accumulates evidence, the particle will move on a plane perpendicular to *t*_2_ = 0, since *t*_2_ remains constant while attending to item 1 (Fig. 2C, first column). During this time, evidence about the true values of both items will be accumulated, but information regarding item 2 will be significantly noisier (as controlled by *γ*). Depending on the evidence accumulated regarding both items, the particle may hit the boundary for âĂIJchoose 1âĂi, âĂIJchoose 2âĂi, or âĂIJswitch (attention)âĂi. Assume the particle hits the âĂIJswitchâĂi boundary, indicating that the model is not confident enough to make a decision after the initial fixation to item 1. Now, the model is attending to item 2, and the policy space switches accordingly (*y* = 2). The particle, starting from where it left off, will now move on a plane perpendicular to the *t*_1_ axis (Fig. 2C, second column). This process is repeated until the particle hits a decision boundary (Fig. 2C, third column). Importantly, these shifts in attention are endogenously generated by the model as a part of the optimal decision strategy — it exploits its ability to control how much information it receives about either item’s value.

The optimal policy space shows some notable properties. As expected, the âĂIJswitchâĂi region in a given policy space is always encompassed in the âĂIJaccumulateâĂi region of the other policy space, indicating that the model never switches attention or makes a decision immediately after an attention switch. Furthermore, the decision boundaries in 3D space approach each other over time, consistent with previous work that showed a collapsing 2D boundary for optimal value-based decisions without attention [42]. The collapsing bound reflects the modelâĂŹs uncertainty regarding the difficulty of the decision task [17]. In our case, this difficulty depends on how different the true item values are, as items of very different values are easier to distinguish than those of similar value. If the difficulty is known within and across choices, the boundaries will not collapse over time, and their (fixed) distance will reflect the difficulty of the choice. However, since the difficulty of individual choices varies and is a priori unknown to the decision maker in our task, the decision boundary collapses so that the model minimizes wasting time on a choice that is potentially too difficult.

The contour of the optimal policy boundaries changes in intuitive ways as different parameters of the model are adjusted (Fig. S1). Increasing the switch cost *c_s_* leads to a smaller policy space for the âĂIJswitchâĂi behavior, since there is an increased cost for switching attention. Similarly, decreasing the inattention noise by increasing *γ* leads to a smaller âĂIJswitchâĂI space because the model can obtain more reliable information from the unattended item, reducing the necessity to switch attention. Increasing the noisiness of evidence accumulation (*σ*^2^) causes an overall shrinkage of the evidence accumulation space. This allows the model to reach a decision boundary more quickly under a relatively higher degree of uncertainty, given that evidence accumulation is less reliable but equally costly. Similarly, increasing the cost of accumulating evidence (*c*) leads to a smaller accumulation space, so that the model minimizes paying a high cost for evidence accumulation.

### The optimal policy replicates human behavior

To assess if the optimal policy features the same decision-making characteristics as human decision makers, we used it to simulate behavior in a task analogous to the value-based decision task performed by humans in Krajbich et al (2010) [25]. Briefly, in this task, participants first rated their preference of different snack items on a scale of 1 to 10. Then, they were presented with pairs of different snacks and instructed to choose the preferred item. While they deliberate on their choice, the participants’ eye movements were tracked and the fixation duration to each item was used as a proxy for visual attention.

We simulated decision-making behavior using value distributions similar to those used in the human experiment (see Methods), and found that the model behavior qualitatively reproduce essential features of human choice behavior (Fig. 3). As expected in value-based decisions, a larger value difference among the compared items made it more likely for the model to choose the higher-valued item (Fig. 3A; *t*(38) = 105.7, *p* < 0.001). Furthermore, the model’s mean response time decreased with increasing value difference, indicating that less time was spent on trials that were easier (Fig. 3B; *t*(38) = – 11.1, *p* < 0.001). The model also made less attentional switches for easier trials, indicating that difficult trials required more evidence accumulation from both items, necessitating multiple switches in attention (Fig. 3C; *t*(38) = – 8.10, *p* < 0.001).

**Figure 3:**
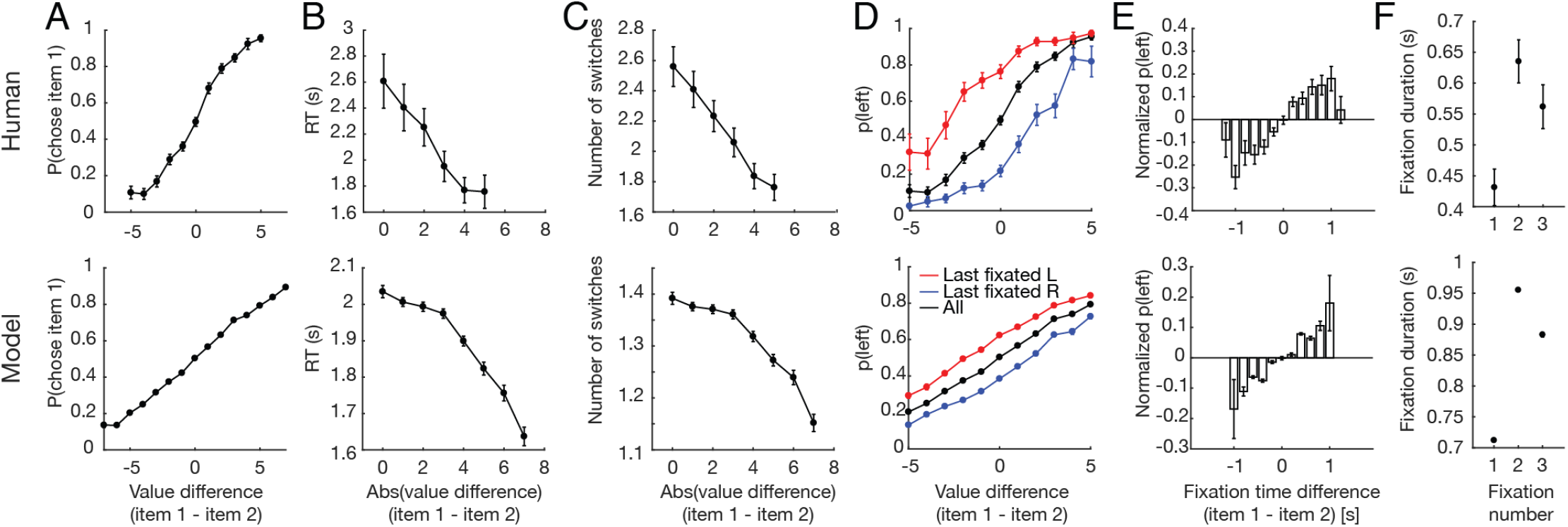
Replication of human behavior by simulated optimal model behavior [25]. (A) Monotonic increase in probability of choosing item 1 as a function of the difference in value between item 1 and 2. (B) Monotonic decrease in response time (RT) as a function of trial difficulty. RT increases with increasing difficulty. (C) Decrease in the number of attention switches as a function of trial difficulty. More switches are made for harder trials. (D) Effect of last fixation location on item preference. The item that was fixated on immediately prior to the decision was more likely to be chosen. (E) Attention’s biasing effect on item preference. The item was more likely to be chosen if it was attended for a longer period of time. (F) Replication of fixation pattern during decision making. Both model and human data showed a fixation pattern where a short initial fixation was followed by a long, then medium-length fixation. Error bars indicate SEM across both human and simulated participants (*N* = 39 for both).

The model also reproduced the biasing effects of fixation on preference seen in humans [25]. An item was more likely to be chosen if it was the last one to be fixated on (Fig. 3D), and if it was viewed for a longer time period (Fig. 3E; *t*(38) = 5.32, *p* < 0.001). Interestingly, the model also replicated a particular fixation pattern seen in humans, where a short first fixation is followed by a significantly longer second fixation, which is followed by a medium-length third fixation (Fig. 3F).

One feature that distinguishes our model from previous attention-based decision models is that attention only modulates the variance of momentary evidence without explicitly down-weighting the value of the unattended item [25, 39]. Therefore, at first glance, preference for the more-attended item is not an obvious feature since our model does not appear to boost its estimated value. However, under the assumption that decision-makers start out with a zero-mean prior, Bayesian belief updating with attention modulation turns out to effectively account for a biasing effect of fixation on the subjective value of items [27]. For instance, consider choosing between two items with equal underlying value. Without an attention-modulated process, the model will accumulate evidence from both items simultaneously, and thus have no preference for one item over the other. However, once attention is introduced and the model attends to item 1 longer than item 2, it will have acquired more evidence about item 1’s value. This will cause item 1 to have a sharper, more certain likelihood function compared to item 2 (Fig. 4A). As posterior value estimates are formed by combining priors and likelihoods in proportion to their associated certainties, the posterior of item 1 will be less biased towards the prior than that of item 2. This leads to a higher subjective value of item 1 compared to that of item 2 even though their true underlying values are equal.

**Figure 4:**
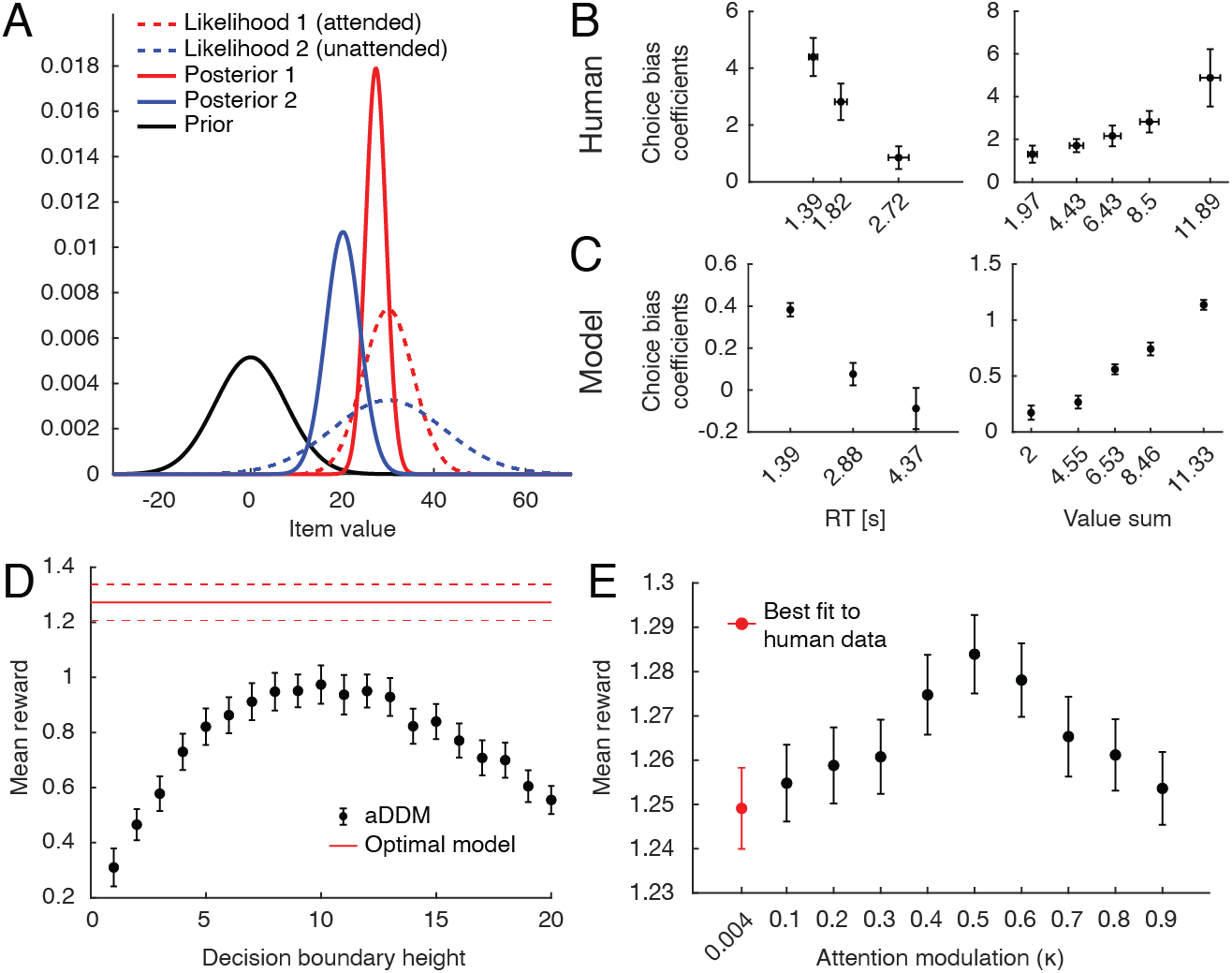
Behavioral predictions from Bayesian value estimation, and further properties of the optimal policy. (A) Bayesian explanation of attention-driven value preference. Attending to one of two equally-valued items for a longer time (red vs. blue) leads to a more certain (i.e., narrower) likelihood and weaker bias of its posterior towards the prior. This leads to a subjectively higher value for longer-attended item. (B,C) Effect of response time (RT; left panel) and sum of the two item values (value sum; right panel) on attention-driven choice bias in humans (B) and the optimal model (C). For (B), the horizontal axis is binned according to number of fixations; the tick marks indicate the mean RT for trials where there were one, two, or three total fixations. For (C), the horizontal axis is binned to contain the same number of trials per bin, where the tick marks indicate the mean value sum for each bin. For (B) and (C), the vertical error bars indicate SEM across participants, and the horizontal error bars indicate SEM across participants of the mean x-values within each bin. (D) Comparing decision performance between the optimal policy and the original aDDM model. Performance of the aDDM was evaluated for different boundary heights (error bars = SEM across simulated participants). Even for the reward-maximizing aDDM boundary height, the optimal model significantly outperformed the aDDM. (E) Decision performance for different degrees of the attention bottleneck (*κ*) while leaving the overall input information unchanged (error bars = SEM across simulated participants). The performance peak for intermediate κ values (i.e., *κ* = 0.5) indicates that allocating similar amounts of attentional resource to both items is beneficial.

This insight leads to additional predictions for how attention-modulated choice bias should vary with certain trial parameters. For instance, the Bayesian account predicts that trials with longer response times should have a weaker choice bias than trials with shorter response times. This is because the difference in fixation times between the two items will decrease over time as the model has more opportunities to switch attention. Both the human and model behavior robustly showed this pattern (Fig. 4B; human, *t*(38) = −3.25, *p* = 0.0024; model, *t*(38) = −32.0, *p* < 0.001). Similarly, choice bias should increase for trials with higher-valued items. In this case, since the evidence distribution is relatively far away from the prior distribution, the posterior distribution is âĂIJpulled awayâĂi from the prior distribution to a greater degree for the attended versus unattended item, leading to greater choice bias. Both human and model data confirmed this behavioral pattern (Fig. 4C; human, *t*(38) = 2.95, *p* = 0.0054; model, *t*(38) = 11.4, *p* < 0.001).

Next, we assessed whether the optimal model outperformed the original attentional drift diffusion model (aDDM) proposed by Krajbich and colleagues (2010), which, despite its simpler structure, could nonetheless provide competitive performance. To ensure a fair comparison, we adjusted the aDDM model parameters (i.e., attentional value discounting and the noise variance) so that the momentary evidence provided to the two models has equivalent signal-to-noise ratios (Supplementary Info). The original aDDM model fixed the decision boundaries at ±1 and subsequently fit model parameters to match behavioral data. Since we were interested in comparing performance, we simulated model behavior using incrementally increasing decision barrier heights, looking for the height that yields the maximum mean reward (Fig. 4D). We found that even for the best-performing decision barrier height, the aDDM model yielded a significantly lower mean reward compared to that of the optimal model (*t*(76) = 3.01, *p* = 0.0027).

Recent advances in artificial intelligence used attentional bottlenecks to regulate information flow with significant associated performance gains [5, 19, 30, 4, 40]. Analogously, attentional bottlenecks might also be beneficial for value-based decision-making. To test this, we asked if paying full attention on a single item at a time confers any advantages over the ability to pay relatively less reliable, but equal attention to multiple options in parallel. To do so, we varied the amount of momentary evidence provided about both the attended and unattended items while keeping the overall amount of evidence fixed. The balance of evidence reliability between the attended and unattended items was controlled by *κ*, such that *κ* = 0 resulted in no evidence about the unattended item, and *κ* = 0.5 resulted in equal momentary evidence about both items (see Supplementary Info). When *κ* = 0.5, switching attention had no effect on the evidence collected about either item. For values *κ* > 0.5, the decision maker collected more evidence about the unattended item compared to the attended item. Therefore, the reliability of evidence from the attended item at *κ* = 0.2 is equal to that of the unattended item at *κ* = 0.8. When tuning model parameters to best match human behavior, we found a low *κ* ≈ 0.004, suggesting that humans tend to allocate the majority of their presumably fixed cognitive resources to the currently attended item. This allows for reliable evidence accumulation for the attended item, but is more likely to necessitate frequent switching of attention.

To investigate whether widening this attention bottleneck leads to changes in decision performance, we simulated model behavior for different values of *κ* (0.1 to 0.9, in 0.1 increments). Interestingly, we found that mean reward from the simulated trials is greatest at *κ* = 0.5 and decreases for more extreme values of *κ*, suggesting that a more even distribution of attentional resources between the two items is beneficial for maximizing reward (*t*(38) = −8.51, *p* < 0.001).

The impact of attention is not unique to value-based decisions. In fact, recent work showed that fixation can bias choices in a perceptual decision-making paradigm [43]. In their task, participants were first shown a target line with a certain orientation, then shown two lines with slightly different orientations. The goal was to choose the line with the closest orientation to the previously shown target. Consistent with results in the value-based decision task, the authors demonstrated that the longer-fixated option was more likely to be chosen. We modified our attention-based optimal policy to perform in such perceptual decisions, in which the goal was to choose the option that is the closest in some quantity to the target, rather than choosing the higher-valued option. Our modified optimal policy was successful at reproducing the attention-driven biases seen in humans (Fig. S2).

## Discussion

In this work, we show that a normative decision-making model with an attentional bottleneck is able to reproduce the choice and fixation patterns of human decision-makers. Unlike previous work that suggested fixation patterns to be independent of the decisionmaking strategy [25, 26], our work shows that they could instead reflect active information gathering through controlling an attentional bottleneck. This interpretation extends previous work on visual attention to the realm of value-based and perceptual decision-making [46, 23, 21, 12, 15].

In contrast to prior models that simply down-weighted the value of the unattended item [25, 26, 39], our model posits that attention enhances the reliability of information [16]. This was inspired by neurophysiological findings demonstrating that visual attention selectively increases the firing rate of neurons tuned to task-relevant stimuli [35], decreases the mean-normalized variance of individual neurons [28, 45], and reduces the correlated variability of neurons at the population level [13, 29, 3]. In essence, selective attention appears to boost the signal-to-noise ratio, or the reliability of information encoded by neuronal signals rather than alter the magnitude of the value encoded by these signals.

When designing our model, we took the simplest possible approach to introduce an attentional bottleneck into normative models of decision-making. Despite this, we were able to capture a wide range of previously observed features of human decisions (Figs. 3 and S2) and confirm new predictions arising from our theory (Fig. 4B&C). When doing so, our aim was not to quantitatively capture all details of human behavior, which might be driven by additional heuristics and features beyond the scope of our model [1, 18]. Instead, we focused on providing a normative explanation for how fixation changes qualitatively interact with human decisions.

Formulating the choice process through Bayesian inference revealed a simple and intuitive explanation for choice biases (Fig. 4A) (see also [27]). This explanation required the decision maker to a-priori believe the items’ values to be lower than they actually are when choosing between appetitive options. The opposite might be true for aversive items, in which decision makers a-priori expect higher values. In this case, our Bayesian framework predicts that choice biases should reverse: less-fixated items should become the preferred choice. This is exactly what has been observed in human decision-makers [2].

Due to the optimal policy’s complexity (Fig. 2), we expect the nervous system to implement it only approximately (e.g., similar to [41] for multi-alternative decisions). Such an approximation has been recently suggested by Callaway and colleagues [10], where they used approaches from rational inattention to approximate optimal decision-making in the presence of an attentional bottleneck. Unlike our work, they assumed that the unattended item is completely ignored, and therefore could not investigate the effect of graded shifts of attentional resources between items (Fig. 4E). In addition, they were unable to replicate the choice bias in binary choices. Nonetheless, despite only approximating the normative strategy, they were able to well-capture many behavioral features of human decisions involving two and three items. Hébert and Woodford have recently addressed a related decision problem in which the amount of attention assigned to either item could be changed continuously over time [22]. While interesting in its own right, such continuous assignment makes it impossible to relate attention to discrete fixation changes, as we do in our work.

We show that adding an attentional bottleneck does not boost performance of our decision-making model (Fig. 4E). Instead, spreading a fixed cognitive reserve evenly between the attended and unattended items maximized performance. Parameters fit to human behavior reveal that humans tend to allocate a large proportion of their cognitive resource toward the attended item, suggesting that the benefits of an attentional bottleneck might lie in other cognitive processes. Indeed, machine learning applied to text translation [5, 19], object recognition [30, 4], and video-game playing [40] benefits from attentional bottlenecks that allow the algorithm to focus resources on specific task subcomponents. For instance, image classification algorithms that extract only the relevant features of an image for high-resolution processing demonstrated improved performance and reduced computational cost compared to those without such attentional features [30]. Similarly, attentional bottlenecks that appear to limit human decision-making performance might have beneficial effects on cognitive domains outside the scope of binary value-based decisions. This is consistent with the idea that the evolutionary advantage of selective attention involves the ability to rapidly fixate on salient features in a cluttered environment, thereby limiting the amount of information that reaches upstream processing and reducing the overall computational burden [24].

## Materials and Methods

Here, we provide an outline of the framework and its results. Detailed derivations are provided in Supplementary Information.

### Attention-modulated decision-making model

Before each trial, *z*_1_ and *z*_2_ are drawn from 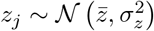. *z*_1_ and *z*_2_ correspond to the value of each item. In each time-step *n* > 0 of duration *δt*, the decision-maker observes noisy samples of each *z_j_*. This momentary evidence is drawn from 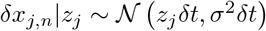 for the attended item *j* = *y_n_*, and 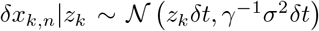 for the unattended item *k* ≠ *y_n_*, where 0 ≤ *γ* ≤ 1 reduces the information provided by this item. The posterior *z_j_* for *j* ∈ {1, 2} after *t* = *Nδt* seconds is found by Bayes’ rule, 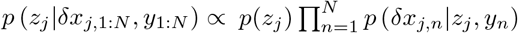, which results in Eq. (2). If *y_n_* ∈ {1, 2} identifies the attended item in each time-step, the attention times in this posterior are given by 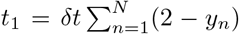 and 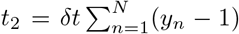. The attention-weighted accumulated evidence is 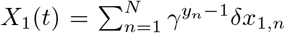 and 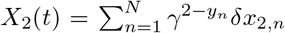, down-weighting the momentary evidence for periods when the item is unattended.

We found the optimal policy by dynamic programming [6, 17], which, at each point in time, chooses the action that promises the larges expected return, including all rewards and costs from that point into the future. Its central component is the value function that specifies this expected return for each value of the sufficient statistics of the task. In our task, the sufficient statistics are the two posterior means, 〈*z_j_*|*X_j_*(*t*), *t*_1_,*t*_2_〉 for *j* ∈ {1, 2}, the two accumulation times, *t*_1_ and *t*_2_, and the currently attended item *y_n_*. The decision maker can choose between four actions at any point in time. The first two are to choose one of the two items, which is expected to yield the corresponding reward, after which the trial ends. The third action is to accumulate evidence for some more time *δt*, which comes at cost *cδt*, and results in more momentary evidence and a corresponding updated posterior. The fourth is to switch attention to the other item 3 – *y_n_*, which comes at cost *c_s_* > 0. As the optimal action is the one that maximizes the expected return, the value for each sufficient statistic is the maximum over the expected returns associated with each action. This leads to the recursive Bellman’s equation that relates values with different sufficient statistics (see SI for details) and reveals the optimal action for each of these sufficient statistics. Due to symmetries in our task, it turns out these optimal actions only depend on the difference in posterior means Δ, rather than each of the individual means (see SI). This allowed us to compute the value function and associated optimal policy in the lower-dimensional (Δ, *t*_1_, *t*_2_, *y*)-space, an example of which is shown in Fig. 2.

The optimal policy was found numerically by backwards induction [42, 9], which assumes that at a large enough *t* = *t*_1_ + *t*_2_, a decision is guaranteed and the expected return equals Δ. We set this time point as *t* = 6*s* based on empirical observations. From this point, we move backwards in small time steps of 0.05*s* and traverse different values of Δ which was also discretized into steps of 0.05. Upon completing this exercise, we are left with a 3-dimensional grid with the axes corresponding to *t*_1_, *t*_2_ and Δ, where the value assigned to each point in space indicates the optimal decision to take for the given set of sufficient statistics. The boundaries between different optimal actions can be visualized as 3-dimensional manifolds (Fig. 2).

### Model simulations

Using the optimal policy, we simulated decisions in a task analogous to the one humans performed in Kracjbich et al., 2010 [25]. On each simulated trial, two items with values *z*_1_ and *z*_2_ are presented. The model attends to one item randomly (*y* ∈ [1, 2]), then starts accumulating noisy evidence and adjusts its behavior across time according to the optimal policy. Since the human data had a total of 39 participants, we simulated the same number of participants (*N* = 39) for the model, but with a larger number of trials. For each simulated participant, trials consisted of all pairwise combinations of values between 0 and 7, iterated 20 times. This yielded a total of 1280 trials per simulated participant.

When computing the optimal policy, there were several free parameters that determined the shape of the decision boundaries. Those parameters included the evidence noise term (*σ*^2^), spread of the prior distribution 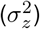, cost of accumulating evidence (*c*), cost of switching attention (*c_s_*), and the relative reliability of evidence accumulation of the attended vs unattended items (*γ*). In order to find a set of parameters that best mimics human behavior, we performed a random search over a large parameter space and simulated behavior using the randomly selected set of parameters [7]. We iterated this process for 2,000,000 sets of parameters and compared the generated behavior to that of humans (see Supplementary Information). After this search process, the parameter set that best replicated human behavior consisted of *c_s_* = 0.0065, *c* = 0.23, *σ*^2^ = 27, 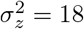, *γ* = 0.004.

### Statistical analysis

The relationship between task variables (e.g., difference in item value) and behavioral measurements (e.g., response time) were assessed by estimating the slope of the relationship for each participant. For instance, to investigate the association between response times and absolute value difference (Fig. 3B), we fit a linear regression within each participant using the absolute value difference and response time for every trial. Statistical testing was performed using one-sample t-tests on the regression coefficients across participants. This procedure was used for statistical testing involving Figs. 3B,C,E, and Figs. 4B,C. To test for a significant peak effect for Fig. 4E, we used the same procedure after subtracting 0.5 to the original *κ* values. To compare performance between the optimal model and the aDDM (Fig. 4D), we first selected the best-performing aDDM model, then performed an independent-samples t-test between the mean rewards from simulated participants from both models.

## Acknowledgments

We would like to thank Ian Krajbich for sharing the behavioral data, and members of the Drugowitsch lab, in particular Anna Kutschireiter and Emma Krause, for feedback on the manuscript. This work was supported by the National Institute of Mental Health (R01MH115554, JD) and the James S. McDonnell Foundation (Scholar Award in Understanding Human Cognition, grant# 220020462, JD).

## References

[1] Acerbi, L., Vijayakumar, S., and Wolpert, D. M. (2014). On the Origins of Suboptimality in Human Probabilistic Inference. PLoS Computational Biology.

[2] Armel, K. C., Beaumel, A., and Rangel, A. (2008). Biasing simple choices by manipulating relative visual attention. Judgment and Decision Making.

[3] Averbeck, B. B., Latham, P. E., and Pouget, A. (2006). Neural correlations, population coding and computation.

[4] Ba, J. L., Mnih, V., and Kavukcuoglu, K. (2015). Multiple object recognition with visual attention. In 3rd International Conference on Learning Representations, ICLR 2015 - Conference Track Proceedings.

[5] Bahdanau, D., Cho, K. H., and Bengio, Y. (2015). Neural machine translation by jointly learning to align and translate. In 3rd International Conference on Learning Representations, ICLR 2015 - Conference Track Proceedings.

[6] Bellman, R. (1952). On the Theory of Dynamic Programming. Proceedings of the National Academy of Sciences.

[7] Bergstra, J. and Bengio, Y. (2012). Random search for hyper-parameter optimization. Journal of Machine Learning Research.

[8] Bertsekas, D. P. (1995). Dynamic programming and optimal control, volume 1. Athena scientific, 4th edition.

[9] Brockwell, A. E. and Kadane, J. B. (2003). A Gridding Method for Bayesian Sequential Decision Problems. Journal of Computational and Graphical Statistics.

[10] Callaway, F., Rangel, A., and Griffiths, T. L. (2020). Fixation patterns in simple choice are consistent with optimal use of cognitive resources. PsyArXiv.

[11] Cavanagh, J. F., Wiecki, T. V., Kochar, A., and Frank, M. J. (2014). Eye tracking and pupillometry are indicators of dissociable latent decision processes. Journal of Experimental Psychology: General.

[12] Chukoskie, L., Snider, J., Mozer, M. C., Krauzlis, R. J., and Sejnowski, T. J. (2013). Learning where to look for a hidden target. Proceedings of the National Academy of Sciences of the United States of America.

[13] Cohen, M. R. and Maunsell, J. H. (2009). Attention improves performance primarily by reducing interneuronal correlations. Nature Neuroscience.

[14] Cohen, M. R. and Maunsell, J. H. (2010). A neuronal population measure of attention predicts behavioral performance on individual trials. Journal of Neuroscience.

[15] Corbetta, M. and Shulman, G. L. (2002). Control of goal-directed and stimulus-driven attention in the brain. Nature Reviews Neuroscience.

[16] Drugowitsch, J., Moreno-Bote, R., and Pouget, A. (2014). Optimal decision-making with time-varying evidence reliability. In Advances in Neural Information Processing Systems.

[17] Drugowitsch, J., Moreno-Bote, R. N., Churchland, A. K., Shadlen, M. N., and Pouget, A. (2012). The cost of accumulating evidence in perceptual decision making. Journal of Neuroscience.

[18] Drugowitsch, J., Wyart, V., Devauchelle, A. D., and Koechlin, E. (2016). Computational Precision of Mental Inference as Critical Source of Human Choice Suboptimality. Neuron.

[19] Gehring, J., Auli, M., Grangier, D., Yarats, D., and Dauphin, Y. N. (2017). Convolutional sequence to sequence learning. In 34th International Conference on Machine Learning, ICML 2017.

[20] Geisler, W. S. and Cormack, L. K. (2012). Models of overt attention. In The Oxford Handbook of Eye Movements.

[21] Hayhoe, M. and Ballard, D. (2005). Eye movements in natural behavior.

[22] Hebert, B. and Woodford, M. (2019). Rational Inattention when decisions take time. Nber Working Paper Series.

[23] Hoppe, D. and Rothkopf, C. A. (2016). Learning rational temporal eye movement strategies. Proceedings of the National Academy of Sciences of the United States of America.

[24] Itti, L. and Koch, C. (2001). Computational modelling of visual attention. Nature Reviews Neuroscience.

[25] Krajbich, I., Armel, C., and Rangel, A. (2010). Visual fixations and the computation and comparison of value in simple choice. Nature Neuroscience.

[26] Krajbich, I. and Rangel, A. (2011). Multialternative drift-diffusion model predicts the relationship between visual fixations and choice in value-based decisions. Proceedings of the National Academy of Sciences of the United States of America.

[27] Li, S. Z. and Ma, W. J. (2019). Valuation as inference: A New Model for the Effects of Fixation on Choice. In Conference on Cognitive Computational Neuroscience.

[28] Mitchell, J. F., Sundberg, K. A., and Reynolds, J. H. (2007). Differential Attention-Dependent Response Modulation across Cell Classes in Macaque Visual Area V4. Neuron.

[29] Mitchell, J. F., Sundberg, K. A., and Reynolds, J. H. (2009). Spatial Attention Decorrelates Intrinsic Activity Fluctuations in Macaque Area V4. Neuron.

[30] Mnih, V., Heess, N., Graves, A., and Kavukcuoglu, K. (2014). Recurrent models of visual attention. In Advances in Neural Information Processing Systems.

[31] Ni, A. M., Ruff, D. A., Alberts, J. J., Symmonds, J., and Cohen, M. R. (2018). Learning and attention reveal a general relationship between population activity and behavior. Science.

[32] Posner, M. I. (1980). Orienting of attention. The Quarterly journal of experimental psychology.

[33] Rangel, A. and Hare, T. (2010). Neural computations associated with goal-directed choice. Current Opinion in Neurobiology.

[34] Ratcliff, R. and McKoon, G. (2008). The diffusion decision model: Theory and data for two-choice decision tasks.

[35] Reynolds, J. H. and Chelazzi, L. (2004). ATTENTIONAL MODULATION OF VISUAL PROCESSING. Annual Review of Neuro-science.

[36] Ruff, D. A., Ni, A. M., and Cohen, M. R. (2018). Cognition as a Window into Neuronal Population Space. Annual Review of Neuroscience.

[37] Shadlen, M. N. N. and Shohamy, D. (2016). Decision Making and Sequential Sampling from Memory. Neuron.

[38] Shimojo, S., Simion, C., Shimojo, E., and Scheier, C. (2003). Gaze bias both reflects and influences preference. Nature Neuroscience.

[39] Song, M., Wang, X., Zhang, H., and Li, J. (2019). Proactive information sampling in value-based decision-making: Deciding when and where to saccade. Frontiers in Human Neuroscience.

[40] Sorokin, I., Seleznev, A., Pavlov, M., Fedorov, A., and Ignateva, A. (2015). Deep attention recurrent Q-network. arXivpreprint arXiv:1512.01693.

[41] Tajima, S., Drugowitsch, J., Patel, N., and Pouget, A. (2019). Optimal policy for multi-alternative decisions. Nature Neuroscience.

[42] Tajima, S., Drugowitsch, J., and Pouget, A. (2016). Optimal policy for value-based decision-making. Nature Communications.

[43] Tavares, G., Perona, P., and Rangel, A. (2017). The attentional Drift Diffusion Model of simple perceptual decision-making. Frontiers in Neuroscience.

[44] Wang, L. and Krauzlis, R. J. (2018). Visual Selective Attention in Mice. Current Biology.

[45] Wittig, J. H., Jang, A. I., Cocjin, J. B., Inati, S. K., and Zaghloul, K. A. (2018). Attention improves memory by suppressing spiking-neuron activity in the human anterior temporal lobe. Nature Neuroscience.

[46] Yang, S. C. H., Lengyel, M., and Wolpert, D. M. (2016). Active sensing in the categorization of visual patterns. eLife.

