## Supplementary Information for "Optimal policy for attention-modulated decisions explains human fixation behavior"

#### Supplementary Text

Here we describe in more detail the derivations of our results, and specifics of the simulations presented in the main text. Of note, we sometimes use $x|y \sim p(y)$ to specify the conditional density $p(x|y)$. Furthermore, $\mathcal{N}\left(\mu, \sigma^2\right)$ denotes a Gaussian with mean $\mu$ and variance $\sigma^2$.

#### 1 Task setup

We assume two latent states $z_j$, $j \in \{1, 2\}$, (here, the true item values) that are before each choice trial drawn from their Gaussian prior, $z_j \sim \mathcal{N}(\bar{z}_j \sigma_z^2)$, with mean $\bar{z}_j$ and variance $\sigma_z^2$. Throughout the text, we will assume $\bar{z}_1 = \bar{z}_2$, to indicate that there is no a-priori preference of one item over the other. The decision maker doesn't observe the latent states, but instead, in each time step of size $\delta t$, observes noisy evidence about both $z_j$'s. Let us assume that, in the $n$th such time step, the decision maker attends to item $y_n \in \{1, 2\}$. Then, they simultaneously observe $\delta x_1$ and $\delta x_2$, distributed as

$$\delta x_{j,n} | y_n, z_j \sim \mathcal{N}\left(z_j \delta t, \frac{1}{\gamma^{|j-y_n|}} \sigma^2 \delta t\right), \tag{1}$$

where we have defined the attention modulation parameter $\gamma$, bounded by $0 \leq \gamma \leq 1$. For the attended item $j = y_n$, we have $|j - y_n| = 0$, such that the momentary evidence $\delta x_{j,n}$ has variance $\sigma^2 \delta t$, independent of $\gamma$. For the unattended item, in contrast, $|j - y_n| = 1$, such that the momentary evidence $\delta x_{j,n}$ has a potentially increased variance $\sigma^2 \delta t / \gamma$, which, for $\gamma < 1$, lowers the information about the underlying latent state. Below we will derive the posterior $z_j$'s, given the stream of momentary evidences $[\delta x_{1,1}, \delta x_{2,1}], [\delta x_{1,2}, \delta x_{2,2}], \dots$, and the attention sequence $y_1, y_2, \dots$. The mean and variance of the posterior distributions represent the decision maker's belief of the items' true values given all available evidence.

While the posterior estimates provide information about value, it does not tell the decision maker when to stop accumulating information, or when to switch their attention. To address these questions, we need to specify the costs and rewards associated with these behaviors. For value-based decisions, we assume that the reward for choosing item $j$ is the latent state $z_j$ (i.e., the true value) associated with the item. Furthermore, we assume that accumulating evidence comes at cost $c$ per second, or cost $c\delta t$ per time step. The decision maker can only ever attend to one item, and switching attention to the other item comes at cost $c_s$ which may be composed of a pure attention switch cost, as well as a loss of time that might introduce an additional cost. As each attention switch introduces both costs, we only consider them in combination without loss of generality.

The overall aim of the decision maker is to maximize the total expected return, which consists of the expected value of the chosen item minus the total cost of accumulating evidence and attention switches. We address this maximization problem by finding the optimal policy that, based on the observed evidence, determines when to switch attention, when to accumulate more evidence, and when to commit to a choice. We initially focus on maximizing the expected return in a single, isolated choice, and will later show that this yields qualitatively similar policies as when embedding this choice into a longer sequence of comparable choices.

#### 2 Bayes-optimal evidence accumulation

##### 2.1 Deriving the posterior $z_1$ and $z_2$

To find the posterior over $z_1$ after having accumulated evidence $x_{1,1:N} \equiv x_{1,1}, \ldots, x_{1,N}$ for some fixed amount of time $t = N\delta t$ while paying attention to items $y_{1:N} \equiv y_1, \ldots y_N$, we employ Bayes' rule,

$$\begin{aligned} p\left(z_1|\delta x_{1,1:N}, y_{1:N}\right) &\propto_{z_1} p(z_1) \prod_{n=1}^{N} p\left(\delta x_{1,n}|z_1, y_n\right) \\ &= \mathcal{N}\left(z_1|\bar{z}_1, \sigma_z^2\right) \prod_{n=1}^{N} \mathcal{N}\left(\delta x_{1,n}|z\delta t, \frac{\sigma^2}{\gamma^{|1-y_n|}}\delta t\right) \\ &\propto_{z_1} \mathcal{N}\left(z_1|\frac{\bar{z}_1\sigma^2\sigma_z^{-2} + X_1(t)}{\sigma^2\sigma_z^{-2} + t_1 + \gamma t_2}, \frac{\sigma^2}{\sigma^2\sigma_z^{-2} + t_1 + \gamma t_2}\right), \end{aligned} \tag{2}$$

where we have defined $X_1(t) = \sum_{n=1}^{N} \gamma^{|1-y_n|}\delta x_{1,n}$ as the sum of all attention-weighted momentary evidence up to time t, and $t_j = t - \delta t \sum_{n=1}^{N} |j - y_n|$ as the total time that item $j$ has been attended. Note that, for time periods in which item 2 is attended to, (i.e., when $y_n = 2$), the momentary evidence is down-weighted by $\gamma$. With $\delta t \to 0$, the process becomes continuous in time, such that $X_1(t)$ becomes the integrated momentary evidence, but the above posterior still holds.

Following a similar derivation, the posterior belief about $z_2$ results in

$$p\left(z_2|\delta x_{2,1:N}, y_{1:N}\right) = \mathcal{N}\left(z_2|\frac{\bar{z}_2\sigma^2\sigma_z^{-2} + X_2(t)}{\sigma^2\sigma_z^{-2} + \gamma t_1 + t_2}, \frac{\sigma^2}{\sigma^2\sigma_z^{-2} + \gamma t_1 + t_2}\right) \tag{3}$$

where $X_2(t) = \sum_{n=1}^{N} \gamma^{|2-y_n|}\delta x_{2,n}$. As the decision maker acquires momentary evidence independently for both items, the two posteriors are independent of each other, that is $p\left(z_1, z_2|\delta x_{1,1:N}, \delta x_{2,1:N}, y_{1:N}\right) = p\left(z_1|\delta x_{1,1:N}, y_{1:N}\right) p\left(z_2|\delta x_{2,1:N}, y_{1:N}\right)$.

##### 2.2 The expected reward process

At each point in time, the decision maker must decide whether it's worth accumulating more evidence versus choosing an item. To do so, they need to predict how the mean estimated reward for each option might evolve if they accumulated more evidence. In this section we derive the stochastic process that describes this evolution for item 1. The same principles will apply for item 2.

Assume that having accumulated evidence until time $t = N\delta t$, the current expected reward for item 1 is given by $\hat{r}_1(t)$, where $\hat{r}_1(t) = \langle z_1|\delta x_{1,1:N}, y_{1:N}\rangle$ is the mean of the above posterior, Eq. (2). The decision-maker's prediction of how the expected reward might evolve after accumulating additional evidence for $\delta t$ is found by the marginalization,

$$\begin{aligned} &p\left(\hat{r}_1(t+\delta t)|\hat{r}_1(t), t_1, t_2, y_{N+1}\right) \\ &\quad = \iint p\left(\hat{r}_1(t+\delta t)|\hat{r}_1(t), \delta x_{1,N+1}, t_1, t_2, y_{N+1}\right) p\left(\delta x_{1,N+1}|z_1, y_{N+1}\right) p\left(z_1|\hat{r}_1(t), t_1, t_2\right) \mathrm{d}\delta x_{1,N+1} \mathrm{d}z_1. \end{aligned} \tag{4}$$

As the last term in the above integral shows, $\hat{r}(t)$, $t_1$ and $t_2$ fully determine the posterior $z_1$ at time $t$. We can use this posterior to predict the value of the next momentary evidence $\delta x_{1,N+1}|z_1$. This, in turn, allows us to predict $\hat{r}_1(t+\delta t)$. As all involved densities are either deterministic or Gaussian, the resulting posterior will be Gaussian as well. Thus, rather than performing the integrals explicitly, we will find the final posterior by tracking the involved means and variances, which in turn completely determine the posterior parameters.

We first marginalize over $\delta x_{1,N+1}$, by expressing $\hat{r}_1(t+\delta t)$ in terms of $\hat{r}(t)$ and $\delta x_{1,N+1}$. To do so, we use Eq. (2) to express $\hat{r}_1(t+\delta t)$ by

$$\hat{r}_1(t+\delta t) = \frac{\bar{z}_1\sigma^2\sigma_z^{-2} + X_1(t) + \gamma^{|y_{N+1}-1|}\delta x_{1,N+1}}{\sigma^2\sigma_z^{-2} + t_1 + \gamma t_2 + \gamma^{|1-y_{N+1}|}\delta t}, \tag{5}$$

where we have used $X_1(t+\delta t) = X_1(t) + \gamma^{|y_{N+1}-1|}\delta x_{1,N+1}$.

Note that, for a given $\delta x_{1,N+1}$, $\hat{r}(t+\delta t)$ is uniquely determined by $\hat{r}(t)$. $\hat{r}(t+\delta t)$ becomes a random variable once we acknowledge that, for any $z_1$, $\delta x_{1,N+1}$ is given by Eq. (1), which we can write as $\delta x_{1,N+1} = z_1\delta t + \sqrt{\sigma^2\gamma^{-|1-y_{N+1}|}\delta t}\eta_x$, where $\eta_x \sim \mathcal{N}(0,1)$. Substituting this expression into $\hat{r}_1(t+\delta t)$, and using Eq. (2) to re-express $X_1(t)$ as $X_1(t) = \hat{r}_1(t)\left(\sigma^2\sigma_z^{-2} + t_1 + \gamma t_2\right) - \bar{z}_1\sigma^2\sigma_z^{-2}$, results in

$$\hat{r}_1(t+\delta t) = \frac{\hat{r}_1(t)\left(\sigma^2\sigma_z^{-2} + t_1 + \gamma t_2\right) + \gamma^{|1-y_{N+1}|}z_1\delta t + \sqrt{\sigma^2\gamma^{|1-y_{N+1}|}\delta t}\eta_x}{\sigma^2\sigma_z^{-2} + t_1 + \gamma t_2 + \gamma^{|1-y_{N+1}|}\delta t}. \tag{6}$$

The second marginalization over $z_1$ is found by noting the distribution of $z_1$ is given by Eq. (2), which can be written as

$$z_1 = \hat{r}_1(t) + \sqrt{\frac{\sigma^2}{\sigma^2\sigma_z^{-2} + t_1 + \gamma t_2}}\eta_z, \tag{7}$$

with $\eta_z \sim \mathcal{N}(0,1)$. Substituting this $z_1$ into the above expression for $\hat{r}(t+\delta t)$ results in

$$\hat{r}_1(t+\delta t) = \hat{r}_1(t) + \frac{\sqrt{\sigma^2 \gamma^{|1-y_{N+1}|}\delta t}}{\sigma^2 \sigma_z^{-2} + t_1 + \gamma t_2 + \gamma^{|1-y_{n+1}|}\delta t}\eta_x, \tag{8}$$

where we have dropped the $\eta_z$-dependent term which had a $\delta t$ pre-factor, and thus vanishes with $\delta t \to 0$. Therefore, $\hat{r}_1(t)$ evolves as a martingale,

$$\hat{r}_1(t+\delta t)|\hat{r}_1(t), t_1, t_2, y_{N+1} \sim \mathcal{N}\left(\hat{r}_1(t), \frac{\sigma^2 \gamma^{|1-y_{n+1}|}}{(\sigma^2 \sigma_z^{-2} + t_1 + \gamma t_2 + \gamma^{|1-y_{N+1}|}\delta t)^2}\delta t\right). \tag{9}$$

Using the same approach, the expected future reward for item 2 is given by

$$\hat{r}_2(t+\delta t)|\hat{r}_2(t), t_1, t_2, y_{N+1} \sim \mathcal{N}\left(\hat{r}_2(t), \frac{\sigma^2 \gamma^{|2-y_{N+1}|}}{(\sigma^2 \sigma_z^{-2} + \gamma t_1 + t_2 + \gamma^{|2-y_{n+1}|}\delta t)^2}\delta t\right). \tag{10}$$

#### 2.3 The expected reward difference process

In a later section, we will reduce the dimensionality of the optimal policy space by using the expected reward difference rather than each of the of the expected rewards separately. To do so, we define this difference by

$$\Delta(t) = \frac{\hat{r}_1(t) - \hat{r}_2(t)}{2}. \tag{11}$$

As for $\hat{r}_1(t)$ and $\hat{r}_2(t)$, we are interested in how $\Delta(t)$ evolves over time.

To find $\Delta(t+\delta t)|\Delta(t), t_1, t_2, y_{N+1}$ we can use

$$p\left(\Delta(t+\delta t)|\Delta(t), t_1, t_2, y_{N+1}\right) = p\left(\Delta(t+\delta t) = \frac{\hat{r}_1(t+\delta t) - \hat{r}_2(t+\delta t)}{2} | \Delta(t) = \frac{\hat{r}_1(t) - \hat{r}_2(t)}{2}, t_1, t_2, y_{N+1}\right). \tag{12}$$

As the decision-maker receives independent momentary evidence for each item, $\hat{r}_1(t)$ and $\hat{r}_2(t)$ are independent when conditioned on $t_1$, $t_2$ and $y_{1:N}$. Thus, so are their time-evolutions, $\hat{r}_1(t+\delta t)|\hat{r}_1(t), \dots$ and $\hat{r}_2(t+\delta t)|\hat{r}_2(t), \dots$. With this, we can show that

$$\Delta(t+\delta t)|\Delta(t), t_1, t_2, y_{N+1} \sim \mathcal{N}\left(\Delta(t), \frac{\sigma^2 \delta t}{4}\left(\frac{\gamma^{|1-y_{N+1}|}}{\left(\sigma^2 \sigma_z^{-2} + t_1 + \gamma t_2 + \gamma^{|1-y_{N+1}|}\delta t\right)^2} + \frac{\gamma^{|2-y_{N+1}|}}{\left(\sigma^2 \sigma_z^{-2} + \gamma t_1 + t_2 + \gamma^{|2-y_{N+1}|}\delta t\right)^2}\right)\right). \tag{13}$$

Unsurprisingly, $\Delta(t)$ is again a martingale.

### 3 Optimal decision policy

We find the optimal decision policy by dynamic programming [1, 3]. A central concept in dynamic programming is the *value function* $V(\cdot)$, which, at any point in time during a decision, returns the *expected return*, which encompasses all expected rewards and costs from that point onwards into the future when following the optimal decision policy. Bellman's equation links value functions across consecutive times, and allows finding this optimal decision policy recursively. In what follows, we first focus on Bellman's equation for single, isolated choices. After that, we show how to extend the same approach to find the optimal policy for long sequences of consecutive choices.

#### 3.1 Single, isolated choice

For a single, isolated choice, accumulating evidence comes at cost $c$ per second. Switching attention comes at cost $c_s$. The expected reward for choosing item $j$ is $\hat{r}_j(t)$, and is given by the mean of Eqs. (2) and (3) for $j=1$ and $j=2$, respectively.

To find the value function, let us assume that we have accumulated evidence for some time $t = t_1 + t_2$, expect rewards $\hat{r}_1(t)$ and $\hat{r}_2(t)$, and are paying attention to item $y \in \{1,2\}$. These statistics fully describe the evidence accumulation state, and thus fully parameterize the value function $V_y(\hat{r}_1, \hat{r}_2, t_1, t_2)$. Here we use $y$ as a subscript rather than an argument to $V(\cdot)$ to indicate that $y$ can only take one of two values, $y \in \{1,2\}$. At this point, we can choose among four actions. We can either immediately choose item 1, immediately choose item 2, accumulate more evidence without switching attention, or switch attention to the other item, $3-y$. The expected return for choosing immediately is either $\hat{r}_1(t)$ or $\hat{r}_2(t)$, depending on the choice. Accumulating more evidence for some time $\delta t$ results in cost $c\delta t$, and changes in the expected rewards according to $\hat{r}_j(t+\delta t)|\hat{r}_j(t), t_1, t_2, y$, as given by Eqs. (9) and (10). Therefore, the expected return for accumulating more evidence is given by

$$-c\delta t + \langle V_y\left(\hat{r}_1(t+\delta t), \hat{r}_2(t+\delta t), t_1 + |2-y|\delta t, t_2 + |1-y|\delta t\right)|\hat{r}_1, \hat{r}_2, t_1, t_2, y\rangle, \tag{14}$$

where the expectation is over the time-evolution of $\hat{r}_1$ and $\hat{r}_2$, and $t_1 + |2-y|\delta t$ and $t_2 + |1-y|\delta t$ ensures that only the $t_y$ associated with the currently attended item is increased by $\delta t$. Lastly, switching attention comes at cost $c_s$, but does not otherwise impact reward expectations, such that the expected return associated with this action is

$$-c_s + V_{3-y}\left(\hat{r}_1, \hat{r}_2, t_1, t_2\right), \tag{15}$$

where the use of $V_{3-y}(\cdot)$ implements that, after an attention switch, item $3-y$ will be the attended item.

By the Bellman optimality principle [1], the best action at any point in time is the one that maximizes the expected return. Combining the expected returns associated with each possible action results in Bellman's equation

$$V_y\left(\hat{r}_1, \hat{r}_2, t_1, t_2\right) = \max \left\{ \begin{array}{c} \hat{r}_1, \hat{r}_2, \\ \left\langle V_y\left(\hat{r}_1(t+\delta t), \hat{r}_2(t+\delta t), t_1 + |2-y|\delta t, t_2 + |1-y|\delta t\right) | \hat{r}_1, \hat{r}_2, t_1, t_2, y \right\rangle - c\delta t, \\ V_{3-y}\left(\hat{r}_1, \hat{r}_2, t_1, t_2\right) - c_s \end{array} \right\}. \tag{16}$$

Solving this equation yields the optimal policy for any combination of $\hat{r}_1$, $\hat{r}_2$, $t_1$, $t_2$ and $y$ by picking the action that maximizes the associated expected return, that is, the term that maximizes the left-hand side of the above equation. The optimal decision boundaries that separate the $(\hat{r}_1, \hat{r}_2, t_1, t_2, y)$-space into regions where different actions are optimal lie at manifolds in which two actions yield the same expected return. For example, the decision boundary at which it becomes best to choose item 1 after having accumulated more evidence is the manifold at which

$$V_y\left(\hat{r}_1, \hat{r}_2, t_1, t_2\right) = \hat{r}_1 = \left\langle V_y\left(\hat{r}_1(t+\delta t), \hat{r}_2(t+\delta t), t_1 + |2-y|\delta t, t_2 + |1-y|\delta t\right) | \hat{r}_1, \hat{r}_2, t_1, t_2, y \right\rangle - c\delta t. \tag{17}$$

In Section 6 we describe how we found these boundaries numerically.

Formulated so far, the value function is five-dimensional, with four continuous ($\hat{r}_1$, $\hat{r}_2$, $t_1$, and $t_2$) and one discrete ($y$) dimension. It turns out that it is possible to remove one of the dimensions without changing the associated policy by focusing on the expected reward difference $\Delta(t)$, Eq. (11), rather than the individual expected rewards. To show this, we jump ahead and use the value function property $V_y\left(\hat{r}_1, \hat{r}_2, t_1, t_2\right) + C = V_y\left(\hat{r}_1 + C, \hat{r}_2 + C, t_1, t_2\right)$ for any scalar $C$, that we will confirm in Sec. 5. Next, we define the value function on expected reward differences by

$$\bar{V}_y(\Delta, t_1, t_2) = V_y(\hat{r}_1, \hat{r}_2, t_1, t_2) - \frac{\hat{r}_1 + \hat{r}_2}{2} = V_y(\Delta, -\Delta, t_1, t_2). \tag{18}$$

Applying this mapping to Eq. (16) leads to Bellman's equation

$$\bar{V}_y\left(\Delta, t_1, t_2\right) = \max \left\{ \begin{array}{c} \Delta, -\Delta, \\ \left\langle \bar{V}_y\left(\Delta(t+\delta t), t_1 + |2-y|\delta t, t_2 + |1-y|\delta t\right) | \Delta, t_1, t_2, y \right\rangle - c\delta t, \\ \bar{V}_{3-y}\left(\Delta, t_1, t_2\right) - c_s \end{array} \right\}, \tag{19}$$

which is now defined over a four-dimensional rather than a five-dimensional space while yielding the same optimal policy. This also confirms that optimal decision-making doesn't require tracking individual expected rewards, but only their difference.

#### 3.2 Sequence of consecutive choices

So far we have focused on the optimal policy for a single isolated choice. Let us now demonstrate that this policy does not qualitatively change if we move to a long sequence of consecutive choices. To do so, we assume that each choice is followed by an inter-trial interval $t_i$ after which the latent $z_1$ and $z_2$ are re-drawn from the prior, and evidence accumulation starts anew. As the expected return considers all expected future rewards, it would grow without bounds for a possibly infinite sequence of choices. Thus, rather than using the value function, we move to using the average-adjusted value function, $\tilde{V}$, which, for each passed time $\delta t$, subtracts $\rho\delta t$, where $\rho$ is the average reward rate [6]. This way, the value tells us if we are performing better or worse than on average, and is thus bounded.

Introducing the reward rate as an additional time cost requires the following changes. First, the average-adjusted expected return for immediate choices becomes $\hat{r}_j(t) - t_i\rho + \tilde{V}_y\left(\bar{z}_1, \bar{z}_2, 0, 0\right)$, where $-t_i\rho$ accounts for the inter-trial interval, and $\tilde{V}_y\left(\bar{z}_1, \bar{z}_2, 0, 0\right)$ is the average-adjusted value at the beginning of the next choice, where $\hat{r}_j = \bar{z}_j$, and $t_1 = t_2 = 0$. Due to the symmetry, $\tilde{V}_y\left(\bar{z}_1, \bar{z}_2, 0, 0\right)$ will be the same for both $y = 1$ and $y = 2$, such that we do not need to specify $y$. Second, accumulating evidence for some duration $\delta t$ now comes at cost $(c + \rho)\delta t$. The expected return for switching attention remains unchanged, as we assume attention switches to be instantaneous. If attention switches take time, we would need to additionally penalize this time by $\rho$.

With these changes, Bellman's equation becomes

$$\tilde{V}_y\left(\hat{r}_1, \hat{r}_2, t_1, t_2\right) = \max \left\{ \begin{array}{c} \hat{r}_1 - \rho t_i + \tilde{V}_y\left(\bar{z}_1, \bar{z}_2, 0, 0\right), \hat{r}_2 - \rho t_i + \tilde{V}_y\left(\bar{z}_1, \bar{z}_2, 0, 0\right), \\ \left\langle \tilde{V}_y\left(\hat{r}_1(t+\delta t), \hat{r}_2(t+\delta t), t_1 + |2-y|\delta t, t_2 + |1-y|\delta t\right) | \hat{r}_1, \hat{r}_2, t_1, t_2, y \right\rangle - (c+\rho)\delta t, \\ \tilde{V}_{3-y}\left(\hat{r}_1, \hat{r}_2, t_1, t_2\right) - c_s \end{array} \right\}. \tag{20}$$

The resulting average-adjusted value function is shift-invariant, that is, adding a scalar to this value function for all states does not change the underlying policy [6]. This property allows us to fix the average-adjusted value for one particular state, such that all other average-adjusted values are relative to this state. For mathematical convenience we choose $\tilde{V}_y(\bar{z}_1, \bar{z}_2, 0, 0) = \rho t_i$, resulting in the new Bellman's equation

$$\tilde{V}_y(\hat{r}_1, \hat{r}_2, t_1, t_2) = \max \left\{ \begin{array}{c} \hat{r}_1, \hat{r}_2, \\ \left\langle \tilde{V}_y(\hat{r}_1(t+\delta t), \hat{r}_2(t+\delta t), t_1 + |2-y|\delta t, t_2 + |1-y|\delta t) \,|\hat{r}_1, \hat{r}_2, t_1, t_2, y \right\rangle - (c+\rho)\delta t, \\ \tilde{V}_{3-y}(\hat{r}_1, \hat{r}_2, t_1, t_2) - c_s \end{array} \right\}. \quad (21)$$

Comparing this to Bellman's equation for single, isolated choices, Eq. (16), reveals an increase in the accumulation cost from $c$ to $c+\rho$. Therefore, we can find a set of task parameters for which the optimal policy for single, isolated choices will mimic that for a sequence of consecutive choices. For this reason, we will focus on single, isolate choices, as they will also capture all policy properties that we expect to see for sequences of consecutive choices.

### 4 Optimal decision policy for perceptual decisions

To apply the same principles to perceptual decision-making, we need to re-visit the interpretation of the latent states, $z_1$ and $z_2$. Those could, for example, be the brightness of two dots on a screen, and the decision-maker needs to identify the brighter dot. Anternatively, they might reflect the length of two lines, and the decision maker needs to identify which of the two lines is longer. Either way, the reward is a function of $z_1$, $z_2$, and the decision maker's choice. Therefore, the expected reward for choosing either option can be computed from the posterior $z$'s, Eqs. (2) and (3). Furthermore, these posteriors are fully determined by their means, $\hat{r}_1$, $\hat{r}_2$, and the attention times, $t_1$ and $t_2$. As a consequence, we can formulate the expected reward for choosing item $j$ by the expected reward function $R_j(\hat{r}_1, \hat{r}_2, t_1, t_2)$.

What are the consequences for this change in expected reward for the optimal policy? If we assume the attention-modulated evidence accumulation process to remain unchanged, the only change is that the expected return for choosing item $j$ changes from $\hat{r}_j$ to $R_j(\hat{r}_1, \hat{r}_2, t_1, t_2)$. Therefore, Bellman's equations changes to

$$V_y(\hat{r}_1, \hat{r}_2, t_1, t_2) = \max \left\{ \begin{array}{c} R_1(\hat{r}_1, \hat{r}_2, t_1, t_2), R_2(\hat{r}_1, \hat{r}_2, t_1, t_2), \\ \langle V_y(\hat{r}_1(t+\delta t), \hat{r}_2(t+\delta t), t_1 + |2-y|\delta t, t_2 + |1-y|\delta t) \,|\hat{r}_1, \hat{r}_2, t_1, t_2, y \rangle - c\delta t, \\ V_{3-y}(\hat{r}_1, \hat{r}_2, t_1, t_2) - c_s \end{array} \right\}. \quad (22)$$

The optimal policy follows from Bellman's equation as before.

The above value function can only be turned into one over expected reward differences under certain regularities of $R_1$ and $R_2$, which we will not discuss further at this point. Furthermore, for the above example, we have assumed two sources of perceptual evidence that need to be compared. Alternative tasks (e.g., the random dot motion task) might provide a single source of evidence that needs to be categorized. In this case, the formulation changes slightly (see, for example, [4]), but the principles remain unchanged.

### 5 Properties of the optimal policy

Here, we will demonstrate some interesting properties of the optimal policy, and the associated value function and decision boundaries. To do so, we re-write the value function in its non-recursive form. To do so, let us first define the switch set $\mathcal{T} = \{T_1, \dots, T_M\}$, which determines the switch times from the current time $t$ onwards. Here, $t + T_1$ is the time of the first switch after time $t$, $t + T_1 + T_2$ is the second switch, and so on. A final decision is made at $t + \bar{T}$, where $\bar{T} = \sum_{m=1}^{M} T_m$, after $M-1$ switches with associated cost $(M-1)c_s$. As the optimal policy is the one that optimizes across choices and switch times, the associated value function can be written as

$$V_y(\hat{r}_1, \hat{r}_2, t_1, t_2) = \max_{\mathcal{T}} \left\langle \max\{\hat{r}_1(t+\bar{T}), \hat{r}_2(t+\bar{T})\} - c\bar{T} - (M-1)c_s |\hat{r}_1, \hat{r}_2, t_1, t_2, y \right\rangle, \quad (23)$$

where time expectation is over the time-evolution of $\hat{r}_1(t)$ and $\hat{r}_2(t)$, that also depends on $\mathcal{T}$. In what follows, we first derive the shift-invarance of this time-evolution, and then consider its consequences for the value function, as well as the decision boundaries.

#### 5.1 Shift-invariance and symmetry of the expected reward process

Let us fix some $\mathcal{T}$, some time $t$, and assume that we are currently attending item 1, $y(t) = 1$. Then, by Eq. (9), $\hat{r}_1(t+\bar{T})$ can be written as

$$\hat{r}_1\left(t+\bar{T}\right) = \hat{r}_1(t) + \int_0^{T_1} \frac{\sigma}{\sigma^2\sigma_z^{-2} + (t_1+s_1) + \gamma t_2} \mathrm{d}B_{1,s_1} + \int_0^{T_2} \frac{\sigma\sqrt{\gamma}}{\sigma^2\sigma_z^{-2} + (t_1+T_1) + \gamma(t_2+s_2)} \mathrm{d}B_{1,s_2}$$
$$+ \int_0^{T_3} \frac{\sigma}{\sigma^2\sigma_z^{-2} + (t_1+T_1+s_3) + \gamma(t_2+T_2)} \mathrm{d}B_{1,s_3} + \dots, \quad (24)$$

where the $B_{1,s_j}$'s are white noise processes associated with item 1. This shows that, for any $\mathcal{T}$, the change in $\hat{r}_1$, that is, $\hat{r}_1\left(t+\bar{T}\right)-\hat{r}_1(t)$, is independent of $\hat{r}_1(t)$. Therefore, we can shift $\hat{r}_1(t)$ by any scalar $C$, and cause an associated shift in $\hat{r}_1\left(t+\bar{T}\right)$, that is

$$p\left(\hat{r}\left(t+\bar{T}\right)=R+C|\hat{r}_1(t)=r+C,t_1,t_2,y\right)=p\left(\hat{r}(t+\bar{T})=R|\hat{r}_1(t)=r,t_1,t_2,y\right), \tag{25}$$

As this holds for any choice of $\mathcal{T}$, it holds for all $\mathcal{T}$. A similar argument establishes this property for $\hat{r}_2$.

The above decomposition of the time-evolution of $\hat{r}_1$ furthermore reveals a symmetry between $\hat{r}_1(t+\bar{T})-\hat{r}_1(t)$ and $\hat{r}_2(t+\bar{T})-\hat{r}_2(t)$. In particular, the same decomposition shows that $\hat{r}_1(t+\bar{T})-\hat{r}_1(t)$ equals $\hat{r}_2(t+\bar{T})-\hat{r}_2(t)$ if we flip $t_1$, $t_2$ and $y(t)$. Therefore,

$$p\left(\hat{r}_1\left(t+\bar{T}\right)=R|\hat{r}_1(t)=r,t_1=a,t_2=b,y=j\right)=p\left(\hat{r}_2\left(t+\bar{T}\right)=R|\hat{r}_2(t)=r,t_1=b,t_2=a,y=3-j\right). \tag{26}$$

#### 5.2 Shift-invariance of the value function

The shift-invariance of $\hat{r}_1$ and $\hat{r}_2$ implies a shift-invariance of the value function. To see this, fix some $\mathcal{T}$ and some final choice $j$, in which case the value function according to Eq. (23) becomes

$$V_y\left(\hat{r}_1,\hat{r}_2,t_1,t_2\right)=\left\langle\hat{r}_j(t+\bar{T})|\hat{r}_1,\hat{r}_2\right\rangle-c\bar{T}+(M-1)c_s, \tag{27}$$

where the expectation is implicitly conditional on $t_1$, $t_2$, $y$ and $\mathcal{T}$. Due to the shift-invariance of the time-evolution of $\hat{r}_1$ and $\hat{r}_2$, adding a scalar $C$ to both $\hat{r}_1$ and $\hat{r}_2$ increases the above expectation by the same amount, $\left\langle\hat{r}_j(t+\bar{T})|\hat{r}_1,\hat{r}_2\right\rangle+C$. As a consequence,

$$V_y\left(\hat{r}_1+C,\hat{r}_2+C,t_1,t_2\right)=V_y\left(\hat{r}_1,\hat{r}_2,t_1,t_2\right)+C. \tag{28}$$

As this holds for any choice of $\mathcal{T}$ and $j$, it also holds for the maximum over $\mathcal{T}$ and $j$, and thus for the value function in general.

A similar argument shows that the value function is increasing in both $\hat{r}_1$ and $\hat{r}_2$. To see this, fix $\mathcal{T}$ and $j$ and note that increasing either $\hat{r}_1$ or $\hat{r}_2$ causes the expectation in Eq. (27) to either remain unchanged or to increase to $\left\langle\hat{r}_j(t+\bar{T})|\hat{r}_1,\hat{r}_2\right\rangle+C$. Therefore, for any non-negative $C$,

$$V_y\left(\hat{r}_1,\hat{r}_2,t_1,t_2\right)\leq V_y\left(\hat{r}_1+C,\hat{r}_2,t_1,t_2\right)\leq V_y\left(\hat{r}_1,\hat{r}_2,t_1,t_2\right)+C, \tag{29}$$

$$V_y\left(\hat{r}_1,\hat{r}_2,t_1,t_2\right)\leq V_y\left(\hat{r}_1,\hat{r}_2+C,t_1,t_2\right)\leq V_y\left(\hat{r}_1,\hat{r}_2,t_1,t_2\right)+C. \tag{30}$$

This again holds for any choice of $\mathcal{T}$ and $j$, such that it holds for the value function in general.

For the value function on expected reward differences, $\bar{V}_y(\Delta,t_1,t_2)$, changing both $\hat{r}_1$ and $\hat{r}_2$ by the same amount leaves $\Delta$, and therefore the associated value $\bar{V}_y\left(\Delta,t_1,t_2\right)$, unchanged. In contrast, increasing only $\hat{r}_1$ or $\hat{r}_2$ by $2C$ increases or decreases $\Delta$ by $C$. Thus, we can use $V_y\left(\hat{r}_1,\hat{r}_2,t_1,t_2\right)=\bar{V}_y\left(\Delta,t_1,t_2\right)+\left(\hat{r}_1+\hat{r}_2\right)/2$ from Eq. (18) and substitute it into the two above inequalities to find

$$\bar{V}_y\left(\Delta,t_1,t_2\right)-C\leq\bar{V}_y\left(\Delta\pm C,t_1,t_2\right)\leq\bar{V}_y\left(\Delta,t_1,t_2\right)+C, \tag{31}$$

for some non-negative $C\geq 0$. This shows that $\bar{V}_y\left(\Delta,t_1,t_2\right)$ changes sublinearly with $\Delta$. However, we cannot anymore guarantee an increase or decrease in $\bar{V}_y(\cdot)$, as an increase in $\Delta$ could arise from both an increase in $\hat{r}_1$ or a decrease in $\hat{r}_2$.

#### 5.3 Symmetry of the value function

The symmetry in time-evolution across $\hat{r}_1$ and $\hat{r}_2$ results in a symmetry in the value function. To show this, let us again fix $\mathcal{T}$ and $j$, such that the value function is given by Eq. (27). Then, by Eq. (26), the expectation in the value function becomes $\left\langle\hat{r}_{3-j}(t+\bar{T})|\hat{r}_2,\hat{r}_1\right\rangle$ if we flip $t_1$, $t_2$, and $j$, while leaving the remaining terms of Eq. (27) unchanged. Therefore,

$$V_y\left(\hat{r}_1,\hat{r}_2,t_1,t_2\right)=V_{3-y}\left(\hat{r}_2,\hat{r}_1,t_2,t_2\right). \tag{32}$$

For the value function on expected reward differences, a flip of $\hat{r}_1$ and $\hat{r}_2$ corresponds to a sign change of $\Delta$, such that we have

$$\bar{V}_y\left(\Delta,t_1,t_2\right)=\bar{V}_{3-y}\left(-\Delta,t_2,t_1\right). \tag{33}$$

Both cases show that we are not required to find the value function for both $y=1$ and $y=2$ separately, as knowing one reveals the other by the above symmetry.

#### 5.4 Maximum $|V_1(\cdot)-V_2(\cdot)|$ difference

By Bellman's equation, Eq. (16), it is best to switch attention if the expected return of accumulating evidence equals that of switching attention, that is, if

$$V_y\left(\hat{r}_1,\hat{r}_2,t_1,t_2\right)=\left\langle V_y\left(\hat{r}_1(t+\delta t),\hat{r}_2(t+\delta t),t_1+|2-y|\delta t,t_2+|1-y|\delta t\right)|\hat{r}_1,\hat{r}_2,t_1,t_2,y\right\rangle-c\delta t=V_{3-y}\left(\hat{r}_1,\hat{r}_2,t_1,t_2\right)-c_s. \tag{34}$$

Before that, $V_{3-y}(\hat{r}_1, \hat{r}_2, t_1, t_2) < V_y(\hat{r}_1, \hat{r}_2, t_1, t_2) + c_s$, as otherwise, an attention switch would have already occurred. When it does, we have $V_{3-y}(\hat{r}_1, \hat{r}_2, t_1, t_2) = V_y(\hat{r}_1, \hat{r}_2, t_1, t_2) + c_s$. That is, the attention switch happens if the value of doing so exceeds that for accumulating evidence by the switch cost $c_s$. Therefore, the difference between the value functions $V_1$ and $V_2$ can never be larger than the switch cost, that is

$$|V_1(\hat{r}_1, \hat{r}_2, t_1, t_2) - V_2(\hat{r}_1, \hat{r}_2, t_1, t_2)| \leq c_s. \tag{35}$$

Once their difference equals the switch cost, a switch occurs. It is easy to see that the same property holds for the value function on expected reward differences, leading to

$$\left|\bar{V}_1(\Delta, t_1, t_2) - \bar{V}_2(\Delta, t_1, t_2)\right| \leq c_s. \tag{36}$$

##### 5.5 The decision boundaries are parallel to the diagonal $\hat{r}_1 = \hat{r}_2$

Following the optimal policy, the decision-maker accumulates evidence until $V_y(\hat{r}_1, \hat{r}_2, t_1, t_2) = \max\{\hat{r}_1, \hat{r}_2\}$. For all times before that, $V_y(\hat{r}_1, \hat{r}_2, t_1, t_2) > \max\{\hat{r}_1, \hat{r}_2\}$, as otherwise, a decision is made. Let us first find an expression for the decision boundaries, and then show that these boundaries are parallel to $\hat{r}_1 = \hat{r}_2$. To do so, we will in most of this section fix $t_1$, $t_2$ and $y$, and drop them for notational convenience, that is $V(\hat{r}_1, \hat{r}_2) \equiv V_y(\hat{r}_1, \hat{r}_2, t_1, t_2)$.

First, let us assume $\hat{r}_1 > \hat{r}_2$, such that $\max\{\hat{r}_1, \hat{r}_2\} = \hat{r}_1$, and item 1 would be chosen if an immediate choice is required. Therefore $V(\hat{r}_1, \hat{r}_2) \geq \hat{r}_1$ always, and $V(\hat{r}_1, \hat{r}_2) = \hat{r}_1$ once a decision is made. For a fixed $\hat{r}_1$, the value function is increasing in $\hat{r}_2$, such that reducing $\hat{r}_2$ if $V(\hat{r}_1, \hat{r}_2) > \hat{r}_1$ will at some point lead to $V(\hat{r}_1, \hat{r}_2) = \hat{r}_1$. The optimal decision boundary is the largest $\hat{r}_2$ for which this occurs. Expressed as a function of $\hat{r}_1$, this boundary on $\hat{r}_2$ is thus given by

$$\theta_{1y}(\hat{r}_1, t_1, t_2) = \max\{\hat{r}_2 \leq \hat{r}_1 : V_y(\hat{r}_1, \hat{r}_2, t_1, t_2) = \hat{r}_1\} \tag{37}$$

A similar argument leads to the optimal decision boundary for item 2. In this case, we assume $\hat{r}_2 > \hat{r}_1$, such that $V(\hat{r}_1, \hat{r}_2) \geq \hat{r}_2$ always, and $V(\hat{r}_1, \hat{r}_2) = \hat{r}_2$ once a decision is made. The sublinear growth of the value function in both $\hat{r}_1$ and $\hat{r}_2$ implies that $V(\hat{r}_1, \hat{r}_2)$ grows at most as fast as $\hat{r}_2$, such that there will be some $\hat{r}_2$ at which $V(\hat{r}_1, \hat{r}_2) > \hat{r}_2$ turns into $V(\hat{r}_1, \hat{r}_2) = \hat{r}_2$. The optimal decision boundary is the smallest $\hat{r}_2$ for which this occurs, that is

$$\theta_{2y}(\hat{r}_1, t_1, t_2) = \min\{\hat{r}_2 \geq \hat{r}_1 : V_y(\hat{r}_1, \hat{r}_2, t_1, t_2) = \hat{r}_2\} \tag{38}$$

Note that both boundaries are on $\hat{r}_2$ as a function of $\hat{r}_1$, $t_1$, $t_2$, and $y$.

To show that these boundaries are parallel to the diagonal, we will use the shift-invariance of the value function, leading, for some scalar $C$, to

$$\begin{aligned}
\theta_{1y}(\hat{r}_1, t_1, t_2) + C &= \max\{\hat{r}_2 + C \leq \hat{r}_1 + C : V_y(\hat{r}_1, \hat{r}_2, t_1, t_2) = \hat{r}_1\} \\
&= \max\{\tilde{r}_2 \leq \tilde{r}_1 : V_y(\tilde{r}_1 - C, \tilde{r}_2 - C, t_1, t_2) = \tilde{r}_1 - C\} \\
&= \max\{\tilde{r}_2 \leq \tilde{r}_1 : V_y(\tilde{r}_1, \tilde{r}_2, t_1, t_2) = \tilde{r}_1\} \\
&= \theta_{1y}(\tilde{r}_1, t_1, t_2) \\
&= \theta_{1y}(\hat{r}_1 + C, t_1, t_2),
\end{aligned} \tag{39}$$

where we have used $\tilde{r}_j = \hat{r}_j + C$. This shows that increasing $\hat{r}_1$ by some scalar $C$ shifts the boundary on $\hat{r}_2$ by the same amount. Therefore, the decision boundary for choosing item 1 is parallel to $\hat{r}_1 = \hat{r}_2$.

An analogous argument for $\theta_{2y}(\cdot)$ results in

$$\theta_{2y}(\hat{r}_1, t_1, t_2) + C = \theta_{2y}(\hat{r}_1 + C, t_1, t_2), \tag{40}$$

which showing that the same property holds for the decision boundary for choosing item 2. Overall, this confirms that the decision boundaries only depend on the expected reward difference (that is, the direction orthogonal to $\hat{r}_1 = \hat{r}_2$), confirming that it is sufficient to compute $\bar{V}(\cdot)$ instead of $V(\cdot)$.

#### 6 Simulation details

##### 6.1 Computing the optimal policy

In Section 3, we described the Bellman equation (Eq. (19)) which outputs the expected return given these four parameters: currently attended item ($y$), reward difference ($\Delta$), expected return for accumulating more evidence, and expected return for switching attention. Note that the symmetry of the value function (Section 5) allows us to drop $-\Delta$ from the original Eq. (19). Solving this Bellman equation provides us with a 4-dimensional "policy space" which assigns the optimal action to take at any point in this space defined by the four parameters above.

The solution to the optimal policy can be found numerically by backwards induction [6]. To do so, first we assume some large $t = t_1 + t_2$, where a decision is guaranteed. In this case, $V_y(\Delta, t_1, t_2) = \max\{-\Delta, \Delta\} = |\Delta|$ for both $y = 1$ and $y = 2$. We call this the base case. From this base case, we can move one time step backwards in $t_1$ $(y = 1)$:

$$\bar{V}_1(\Delta, t_1 - \delta t, t_2) = \max \left\{ \begin{array}{c} \Delta, \\ \langle \bar{V}_1(\Delta, t_1, t_2) \, | \Delta, t_1, t_2 \rangle - c\delta t, \\ \bar{V}_2(\Delta, t_1 - \delta t, t_2) - c_s \end{array} \right\}, \quad (41)$$

The second expression in the maximum can be evaluated, since we assume a decision is made at time t. But $\bar{V}_2(\Delta, t_1 - \delta t, t_2) - c_s$, which is the value function for switching attention, is unknown. This unknown value function is given by

$$\bar{V}_2(\Delta, t_1 - \delta t, t_2) = \max \left\{ \begin{array}{c} \Delta, \\ \langle \bar{V}_2(\Delta, t_1 - \delta t, t_2 + \delta t) \, | \Delta, t_1, t_2 \rangle - c\delta t, \\ \bar{V}_1(\Delta, t_1 - \delta t, t_2) - c_s \end{array} \right\}, \quad (42)$$

In this expression, the second term can again be found, but $\bar{V}_1(\Delta, t_1 - \delta t, t_2) - c_s$ is unknown. Looking at the two expressions above, we see that under the parameters $(\Delta, t_1 - \delta t, t_2)$, $V_1 \geq V_2 - c_s$, and $V_2 \geq V_1 - c_s$, which cannot both be true. Therefore, we first assume that $V_1$ is not determined by $V_2 - c_s$, removing the $V_2 - c_s$ term from the maximum. This allows us to find $\bar{V}_1(\Delta, t_1 - \delta t, t_2)$ in Eq. (41). Then, we compute Eq. (42) including the $V_1 - c_s$ term. If we find that $V_2 = V_1 - c_s$, then $V_1 \neq V_2 - c_s$, which means the $V_2 - c_s$ term could not have mattered in Eq. (41), and we are done. If not, we re-compute $V_1$ with the $V_2 - c_s$ term included, and we are done. Therefore, we were able to compute $V_1$ and $V_2$ under the parameters $(\Delta, t_1 - \delta t, t_2)$ using information about $\bar{V}_1(\Delta, t_1, t_2)$ and $\bar{V}_2(\Delta, t_1 - \delta t, t_2 + \delta t)$.

Using the same approach, we can find $V_{1,2}(\Delta, t_1, t_2 - \delta t)$ based on $\bar{V}_1(\Delta, t_1 - \delta t, t_2 + \delta t)$ and $\bar{V}_2(\Delta, t_1, t_2)$. Thus, given that we know $V_y(\Delta, t_1, t_2)$ above a certain $t = t_1 + t_2$, we can move backwards to compute $V_1$ and $V_2$ for $(\Delta, t_1 - \delta t, t_2)$, then $(\Delta, t_1 - 2\delta t, t_2)$, and so on, until $(\Delta, 0, t_2)$ for all relevant values of $\Delta$. Subsequently, we can do the same moving backwards in $t_2$, solving for $V_y(\Delta, t_1, t_2 - \delta t)$, $V_y(\Delta, t_1, t_2 - 2\delta t)$, ..., $V_y(\Delta, t_1, 0)$. Following this, we can continue with the same procedure from $V_y(\Delta, t_1 - \delta t, t_2 - \delta t)$, until we have found $V_{1,2}$ for all combinations of $t_1$ and $t_2$.

In practice, the parameters of the optimal policy space were discretized to allow for tractable computation. We set the large time at which decisions are guaranteed at $t = 6s$, which we determined empirically. Time was discretized into steps of $\delta t = 0.05s$. The item values, and their difference ($\Delta$) were also discretized into steps of 0.05.

Upon completing this exercise, we now have two 3-dimensional optimal policy spaces. The decision-maker's location in this policy space is determined by $t_1$, $t_2$, and $\Delta$. Each point in this space is assigned an optimal action to take (choose item, accumulate more evidence, switch attention) based on which expression was largest in the maximum of the respective Bellman equation. The decision-maker moves between the two policy spaces depending on which item they are attending to ($y \in [1, 2]$).

In order to find the 3-dimensional boundaries that signify a change in optimal action to take, we took slices of the optimal policy space in planes of constant $\Delta$'s. We found the boundary between different optimal policies within each of these slices. We in turn approximated the 3-dimensional contour of the optimal policy boundaries by collating them along the different $\Delta$'s.

#### 6.2 Finding task parameters that best match human behavior

In computing the optimal policy, there were several free parameters that determined the shape of the policy boundaries, thereby affecting the behavior of the optimal model. These parameters included $\sigma^2$, $\sigma_z^2$, $c$, $c_s$, and $\gamma$. Our goal was to find a set of parameters that qualitatively mimic human behavior as best as possible. To do so, we performed a random search over the following parameter values: $c_s \in [0.001, 0.05]$ (steps size 0.001), $c \in [0.01, 0.4]$ (steps size 0.01), $\sigma^2 \in [1, 100]$ (step size 1), $\sigma_z^2 \in [1, 100]$ (step size 1), $\gamma \in [0.001, 0.01]$ (step size 0.001) [2].

To find the best qualitative fit, we simulated behavior from a randomly selected set of parameter values (see next section for simulation procedure). From this simulated behavior, we evaluated the match between human and model behavior by applying the same procedure to each of Figs. 3B, C, E. For each bin for each plot, we subtracted the mean values between the model and human data, then divided this difference by the standard deviation of the human data corresponding to that bin, essentially computing the effect size of the difference in means. We computed the sum of these effect sizes for every bin, which served as a metric for how qualitatively similar the curves were between the model and human data. We performed the same procedure for all three figures, and ranked the sum of the effect sizes for all simulations. We performed simulations for over 2,000,000 random sets of parameter values. The set of parameters for which our model best replicated human behavior according to the above criteria was $c_s = 0.0065$, $c = 0.23$, $\sigma^2 = 27$, $\sigma_z^2 = 18$, $\gamma = 0.004$.

#### 6.3 Simulating decisions with the optimal policy

The optimal policy allowed us to simulate decision making in a task analogous to the one humans performed in Kracjbich et al., 2010 [5]. For a given set of parameters, we first computed the optimal policy. In a simulated trial, two items with values $z_1$ and $z_2$ are presented. At trial onset, the model attends to an item randomly ($y \in [1, 2]$), and starts accumulating noisy evidence centered around the true values. At every time step ($\delta t = 0.05$), the model evaluates $\Delta$ using the mean of the posteriors between the two items (see

Eqs. (2) and (3)). Then, the model performs the optimal action associated with its location in the optimal policy space. If the model makes a decision, then the trial is over. If the model instead accumulates more evidence, then the above procedure is repeated for the next time step. If the model switches attention, it does not obtain further information about either item, but switches attention to the other item. Switching attention allows for more reliable evidence from the now-attended item, and also switches the optimal policy space to the appropriate one (see Figure 2).

To allow for a relatively fair comparison between the model and human data, we simulated the same number of subjects ($N = 39$) for the model, but with a larger number of trials. For each simulated subject, trials were created such that all pairwise combinations of values between 0 and 7 were included, and this was iterated 20 times. This yielded a total of 1280 trials per subject.

#### 6.4 Attention diffusion model

In order compare the decision performance of the optimal model to that of the original attentional drift diffusion model (aDDM) proposed by Krajbich and co1leagues [5], we needed to ensure that neither model had an advantage by receiving more information. We did so by making sure that the signal-to-noise ratios of evidence accumulation of both models were identical. In aDDM, the evidence accumulation evolved according to the following process, in steps of 0.05s (assuming y = 1):

$$v_t = v_{t-1} + d(z_1 - \gamma_k z_2) + \eta_t, \tag{43}$$

where $v_t$ is the relative decision value that represents the subjective value difference between the two items at time $t$, $d$ is a constant that controls the speed of integration (in $ms^{-1}$), $\gamma_k$ controls the biasing effect of attention, and $\eta_t \sim \mathcal{N}\left(0, \sigma^2\right)$ is a normally distributed random variable zero mean and variance $\sigma^2$. Written differently, the difference in the attention-weighted momentary evidence between item 1 and item 2 can be expressed as

$$\begin{aligned} \delta\Delta = d\left(z_1 - \gamma_k z_2\right) + \eta_t &\sim \mathcal{N}\left(d(z_1 - \gamma_k z_2), \sigma^2\right) \\ &\sim \mathcal{N}\left(k(z_1 - \gamma_k z_2)\delta t, \sigma_k^2 \delta t\right), \end{aligned} \tag{44}$$

where $d$ and $\sigma^2$ were replaced by $k\delta t$, and $\sigma_k^2 \delta t$, respectively. Here, the variance term $\sigma_k^2 \delta t$ can be split into two parts, such that the $\delta\Delta$ term can be expressed as

$$\delta\Delta \sim \mathcal{N}\left(z_1 k \delta t, \frac{1}{2}\sigma_k^2 \delta t\right) - \mathcal{N}\left(\gamma_k z_2 k \delta t, \frac{1}{2}\sigma_k^2 \delta t\right). \tag{45}$$

The signal-to-noise ratios (i.e., the ratio of mean over standard deviation) of the two terms in the above equation are $\frac{z_1 k \delta t}{\sqrt{\frac{\delta t}{2}}\sigma_k}$ and $\frac{z_2 k \delta t}{\frac{1}{\gamma_k}\sigma_k \sqrt{\frac{\delta t}{2}}}$, respectively.

Continuing to assume $y = 1$, in the Bayes-optimal model, evidence accumulation evolves according to

$$\begin{aligned} \delta x_1 &\sim \mathcal{N}\left(z_1 \delta t, \sigma_b^2 \delta t\right), \\ \delta x_2 &\sim \mathcal{N}\left(z_2 \delta t, \gamma_b^{-1} \sigma_b^2 \delta t\right). \end{aligned} \tag{46}$$

Therefore, the difference in the attention-weighted momentary evidence between item 1 and item 2 can be expressed as:

$$\begin{aligned} \delta\Delta &\sim \mathcal{N}\left(z_1 \delta t, \sigma_b^2 \delta t\right) - \gamma_b \mathcal{N}\left(z_2 \delta t, \gamma_b^{-1} \sigma_b^2 \delta t\right) \\ &\sim \mathcal{N}\left(z_1 \delta t, \sigma_b^2 \delta t\right) - \mathcal{N}\left(\gamma_b z_2 \delta t, \gamma_b \sigma_b^2 \delta t\right). \end{aligned} \tag{47}$$

The signal-to-noise ratios of the two terms in the above equation are $\frac{z_1 \delta t}{\sqrt{\delta t}\sigma_b}$ and $\frac{z_2 \delta t}{\frac{1}{\sqrt{\gamma_b}}\sigma_b \sqrt{\delta t}}$, respectively.

In order to match the signal-to-noise ratios of the two models, we set equal their corresponding expressions, to find the following relationship between the parameters of the two models:

$$\begin{aligned} k &= 1, \\ \sigma_k^2 &= 2\sigma_b^2, \\ \gamma_k &= \sqrt{\gamma_b}. \end{aligned} \tag{48}$$

Therefore, we simulated the aDDM with model parameters $\gamma_k = \sqrt{\gamma_b}$ and $\sigma_k^2 = 2\sigma_b^2$.

In the original aDDM model, the model parameters were estimated by fitting the model behavior to human behavior after setting a decision threshold at $\pm 1$. Since we adjusted some of the aDDM parameters, we instead iterated through different decision thresholds (1 through 10, in increments of 1) and found the value that maximizes model performance. To keep it consistent with behavioral data, we generated 39 simulated participants that each completed 200 trials where the two item values were drawn from the prior distribution of the optimal policy model, $z_j \sim \mathcal{N}\left(\bar{z}, \sigma_z^2\right)$ using both the optimal model and the aDDM model.

#### 6.5 Adjusting the attention bottleneck

We investigated whether changing the relative amount of attentional resource dedicated to the attended versus unattended item would influence decision-making performance. To do so, we varied the amount of momentary evidence provided about the attended and unattended items while keeping the overall evidence constant. We found the overall evidence from the base model by computing the Fisher information ($I_{base}$) it provides about the respective true item values. This Fisher information is computed as the sum of the reciprocal of the variance from the attended and unattended items, resulting in

$$I_{base} = \frac{1}{\sigma^2} + \frac{1}{\gamma^{-1}\sigma^2} = \frac{1+\gamma}{\sigma^2}. \tag{49}$$

Our goal is to use $\kappa$ $(0 \leq \kappa \leq 1)$ to control the relative attentional resource allocated to the attended versus unattended item, analogous to the $\gamma$ term used in the base model. To do so, we set the variance of the two items as $\sigma_{tot}^2/(1-\kappa)$ and $\sigma_{tot}^2/\kappa$ for the attended and unattended items, respectively, where $\sigma_{tot}^2 = \frac{1}{I_{base}}$ represents the total variance associated with evidence accumulation of both items. This satisfies our requirement of flexibly changing attention allocation while maintaining the Fisher information of the base model,

$$\frac{1-\kappa}{\sigma_{tot}^2} + \frac{\kappa}{\sigma_{tot}^2} = \frac{1}{\sigma_{tot}^2} = I_{base}. \tag{50}$$

To implement this adjusted model, for each value of $\kappa$, we found the associated $\sigma_\kappa^2$ and $\gamma_\kappa$ to replace the $\sigma^2$ and $\gamma$ terms in the base model. To do so, we set the variance of the attended item above equal to that from the base model,

$$\frac{1-\kappa}{\sigma_{tot}^2} = \frac{1}{\sigma_\kappa^2}. \tag{51}$$

Since $\sigma_{tot}^2 = \frac{1}{I_{base}} = \frac{\sigma^2}{1+\gamma}$, we can rearrange the above and solve for $\sigma_k^2$ and $\gamma_\kappa$ to get,

$$\begin{aligned} \sigma_\kappa^2 &= \frac{\sigma_{tot}^2}{1-\kappa}, \\ \gamma_\kappa &= \frac{\kappa}{1-\kappa}. \end{aligned} \tag{52}$$

Using the above $\sigma_\kappa^2$ and $\gamma_\kappa$, we computed the optimal policy and simulated behavior using the same approach as for the base model.

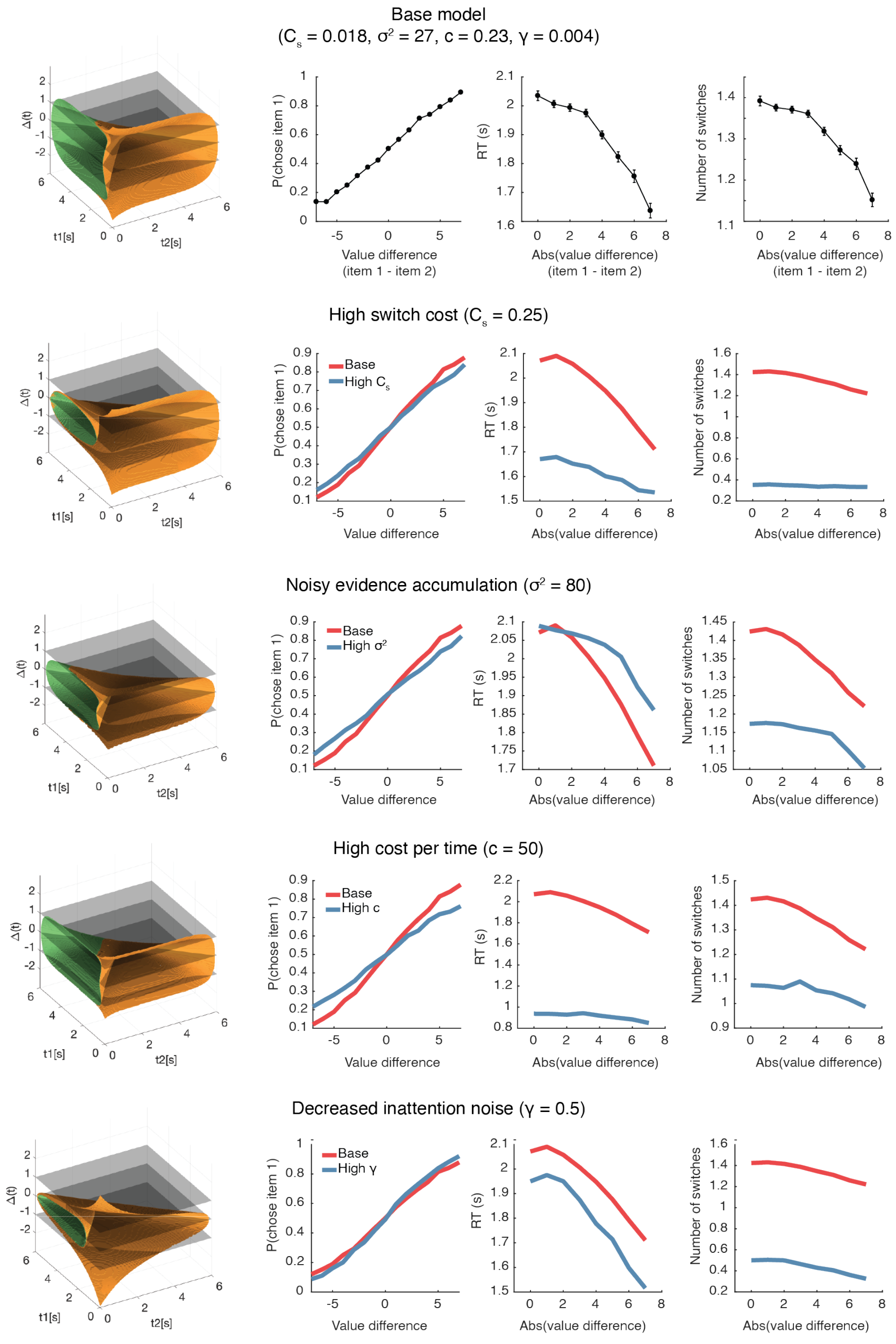


Figure 1: Changes in the optimal policy space and model behavior with adjustments in free model parameters. The optimal policy space and its associated psychometric curves from the base model is shown in the top row. The policy space and psychometric curves corresponding to changes in single free parameters are shown in subsequent rows. In rows 2-4, psychometric curves from he base model on row 1 is shown in red for comparison.

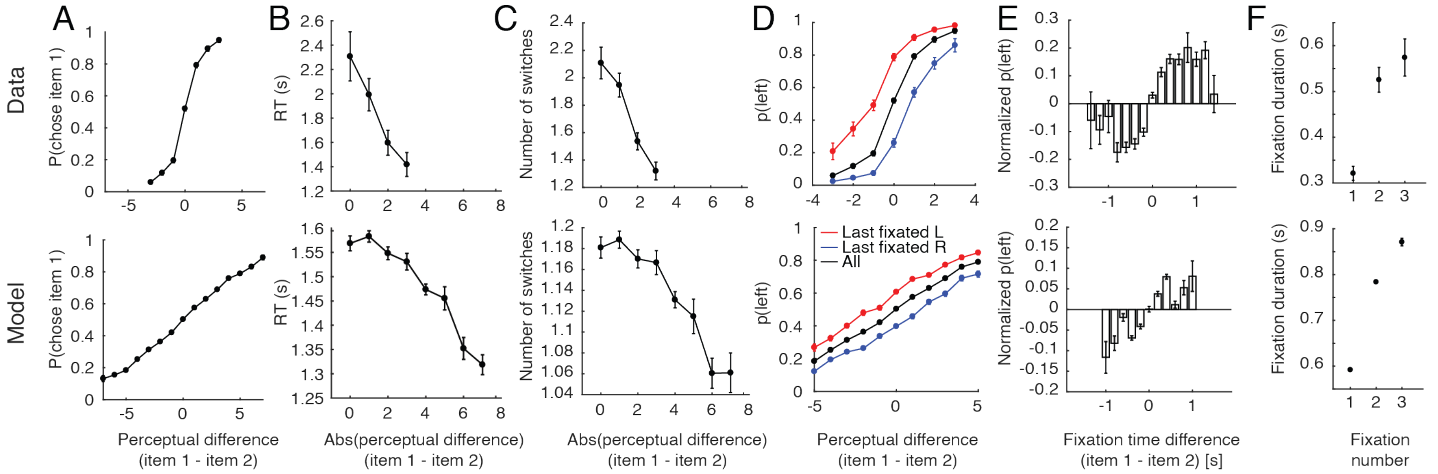


Figure 2: Replication of human behavior by simulated optimal model behavior in a perceptual decision-making task. This task involves choosing the item with a greater degree of a certain a perceptual quality (e.g., brightness of a dot, angle of a line). Therefore, the decision maker is interested in the difference in the perceptual quality between the two items, rather than their difference in value. (A) Monotonic increase in probability of choosing item 1 as a function of the perceptual difference between item 1 and 2. (B) Monotonic decrease in response time (RT) as a function of trial difficulty. (C) Decrease in the number of switches as a function of trial difficulty. (D) Effect of last fixation location on item preference. The item that was fixated on immediately prior to the decision was more likely to be chosen. (E) Attention's biasing effect on item choice. The item was more likely to be chosen if it was attended to for a longer period of time. (F) Replication of fixation pattern during decision making. In the perceptual decision-making task, both model and human data showed increased duration for every subsequent fixation, a notable difference compared to fixation behavior in the value-based task. For (A)-(D), the behavioral data has a smaller range of perceptual difference due to insufficient trials with such large perceptual difference. Error bars indicate SEM across participants for both human and simulated data.

### References

[1] Bellman, R. (1952). On the Theory of Dynamic Programming. *Proceedings of the National Academy of Sciences*.

[2] Bergstra, J. and Bengio, Y. (2012). Random search for hyper-parameter optimization. *Journal of Machine Learning Research*.

[3] Bertsekas, D. P. (1995). *Dynamic programming and optimal control*, volume 1. Athena scientific Belmont, MA.

[4] Drugowitsch, J., Moreno-Bote, R. N., Churchland, A. K., Shadlen, M. N., and Pouget, A. (2012). The cost of accumulating evidence in perceptual decision making. *Journal of Neuroscience*.

[5] Krajbich, I., Armel, C., and Rangel, A. (2010). Visual fixations and the computation and comparison of value in simple choice. *Nature Neuroscience*.

[6] Tajima, S., Drugowitsch, J., and Pouget, A. (2016). Optimal policy for value-based decision-making. *Nature Communications*.
